# Endoplasmic reticulum-mitochondrion disconnection promotes metabolic reprogramming and cystogenesis in polycystic kidney disease

**DOI:** 10.1101/2025.10.31.685870

**Authors:** Biswajit Padhy, Jian Xie, Danish Idrees, Chih-Jen Cheng, Chou-Long Huang

**Affiliations:** Department of Medicine, Division of Nephrology, University of Iowa Carver College of Medicine, Iowa City, Iowa, USA

**Keywords:** ADPKD, Polycystin-1, Polycystin-2, Mitochondria, Endoplasmic reticulum

## Abstract

Mutations in *PKD1* and *PKD2* cause autosomal-dominant polycystic kidney disease (ADPKD), characterized by fluid-filled cysts, aberrant cell proliferation, and widespread genetic and epigenetic remodeling. While mitochondrial dysfunction and metabolic shifts are central to disease progression, the mechanisms linking *PKD* mutations to these changes remain unclear. Here, we demonstrate that ER-mitochondria connectivity was disrupted in *Pkd1- and Pkd2-*deleted mice, preceding cyst formation. This disconnection induces mitochondrial stress, triggering epigenetic remodeling and transcriptional activation of pathways driving proliferation and metabolic reprogramming. Remarkably, restoring *PKD* function in the ER or pharmacologically enhancing ER-mitochondria connection ameliorates mitochondrial dysfunction, epigenetic shifts, and cystogenesis. These findings reveal a critical role for ER-localized PKD in maintaining mitochondrial integrity and transcriptional homeostasis. Mitochondrial dysfunction resulting from ER-mitochondria uncoupling emerges as a key driver of cystogenesis in ADPKD, and correcting this defect may offer a promising therapeutic strategy.

**Significance:** Autosomal dominant polycystic kidney disease (ADPKD) is the most prevalent monogenic cause of kidney failure, marked by fluid-filled cysts, aberrant cell proliferation, metabolic reprogramming, and extensive genetic and epigenetic alterations. The mechanisms by which loss-of-function mutations in *PKD1* and *PKD2* drive disease progression remain poorly understood. Here, we demonstrate that ER-mitochondria contacts are disrupted in *Pkd*-mutant mice prior to cyst formation. This disconnection induces mitochondrial dysfunction and epigenetic remodeling, which in turn promote metabolic reprogramming and cystogenesis. Restoration of *PKD* function in the ER or pharmacological enhancement of ER-mitochondria coupling mitigates these pathological changes. Our findings uncover a critical role for ER-mitochondria crosstalk in suppressing cystogenesis and identify a promising therapeutic target for ADPKD.

## Introduction

Autosomal dominant polycystic kidney disease (ADPKD) is characterized by formation of fluid-filled cysts in the kidney.^1^ Mutations in *PKD1* and *PKD2* coding for polycystin-1 (PC1) and polycystin-2 (PC2), respectively, are responsible for the majority of ADPKD cases. PC1 and PC2 are localized to the primary cilium, cell membrane, and intracellular organelles including the endoplasmic reticulum (ER).^1,2^ How loss of PC1 or PC2 function in cilia and/or other compartments leads to cystogenesis remains elusive.

Metabolic disturbances play important roles in the pathogenesis of ADPKD. *PKD1* mutant cells exhibited enhanced glycolysis and decreased mitochondrial tricarboxylic acid cycle (TCA) reactions and oxidative phosphorylation (OXPHOS).^3^ These metabolic switches in the presence of adequate oxygen supply resemble the Warburg effect observed in tumor cells.^4^ The importance of metabolic reprogramming in ADPKD is supported by studies showing correcting the disturbances ameliorates the disease.^4^ Besides the metabolic switches, mitochondria morphological changes occurs in ADPKD mutant cells.^5^ The mechanism causes these changes is unknown.

We have recently reported that ER-localized PC2 is important in anti-cystogenesis.^6^ Earlier studies have shown ER Ca^2+^ release defects in PC2-deficient cells.^2^ Yet, PC2 is a non-selective cation channel more permeant for K^+^ than Ca^2+^.^7^ We showed that ER-localized PC2 promotes K^+^-Ca^2+^ exchange to facilitate ER Ca^2+^ release.^6^ Expression of an ER-restricted K^+^-permeable channel, trimeric intracellular channel type-B (TricB), corrects ER Ca^2+^ release defect in PC2-null cells and ameliorates cyst formation in *Pkd2*-cKO mice.^6^ ER and mitochondria are two major organelles that occupy large parts of the intracellular space. ER and mitochondria form close contacts inside cells in structures known as mitochondria-associated ER membranes (MAMs) or mitochondria-ER contact sites (MERCS).^8^ The separation distance between ER and mitochondrial membranes at MAM contact sites depend on the type of ER.^9^ The large ribosome constrains the distance between the rough ER and mitochondria to 50-80 nm. The contact between the smooth ER and mitochondria is typically closer to 10-30 nm. The MAM contact between mitochondria and the smooth ER is critical for direct ER-mitochondria Ca^2+^ transfer whereas that between mitochondria and the rough ER is for proteins and lipids transfer.^9^ In renal tubules, a large portion of mitochondrial surface are covered by the ER membranes.

We hypothesize that ER-localized polycystins regulate mitochondrial health and the ER and mitochondria crosstalk is important for anti-cystogenesis. We show that *Pkd1*- and *Pkd2*-cKO kidneys exhibited mitochondrial morphological changes including disruption of MAM contacts. The changes occur early at the pre-cystic stage. Along with the structural changes, mitochondrial respiratory function and genes that regulate mitochondrial biogenesis and dynamics are altered. Cystogenesis, mitochondrial functional and morphological alterations are all reversed by expression of the *TricB* transgene. Mitochondria are not only the powerhouse of cells but also signaling organelles. We further show that mitochondrial dysfunction increases enhancer histone acetylation and expression of *cMyc* and *Cdk7*, a gene crucial for the assembly of super-enhancer complexes for metabolic reprogramming genes.^10^ These mitochondria-driven epigenetic shifts are reversed by the *TricB* transgene. Finally, we show that a pharmacological activator of MAMs function mitigates mitochondrial structural and functional changes, genetic and epigenetic landscape alterations, and cystogenesis in ADPKD mice. Thus, *PKD* mutation-associated defects in ER Ca^2+^ homeostasis disrupt mitochondrial structure and function. Mitochondrial dysfunction fuels epigenetic rewiring amplifying the disease. Enhancing ER-mitochondrial connection may be an effective treatment for the disease.

## Results

### Mitochondrial structural alterations in Pkd2-cKO kidneys and amelioration by correcting ER Ca^2+^ homeostasis defects by TricB expression

We investigated potential causal relationships between mitochondrial dysfunction and cystogenesis examining mitochondrial morphology in *Pkd2*-cKO at the pre-cystic as well as cystic stage. *Pkd2*-cKO kidneys at the pre-cystic stage were similar in size to the control kidneys whereas cystic kidneys were markedly enlarged (Fig. 1, A and B). Blood urea nitrogen (BUN) levels from mice at the cystic (but not the pre-cystic) stage were elevated vs controls (Fig. 1C). We examined the structure and morphology of mitochondria in tubules including proximal (PTs) and distal (DTs) by transmission electron microscopy (TEM). Compared to those in control, mitochondria (marked by blue arrows) in both PTs and DTs of *Pkd2*-cKO kidneys were markedly reduced in number (Fig. 1D; Fig. S1). In addition, mitochondria in the cKO tubules exhibited morphological alterations (from elongated to rounded), decreased cristae density, and were smaller evident by decreased area per mitochondrion (Fig. 1, D-F; Fig. S1). These changes were readily apparent at the pre-cystic stage of cKO kidney. Interestingly, along with the reduced number of mitochondria in cKO tubules, lysosomes (red arrows) were increased (Fig. 1; Fig S1). Others have reported upregulation of the lysosome biogenesis master regulator TFEB and lysosomal expansion in *Pkd*-mutant mutant mice and cell lines.^11^ We have demonstrated the important role of ER Ca^2+^ in anti-cystogenesis and that correcting PC2-dependent ER Ca^2+^ release defects by *TricB* transgene expression reverses cystogenesis in *Pkd2*-cKO.^6^ To test the hypothesis that defects in ER Ca^2+^ homeostasis underlies the mitochondrial alterations, we examined the effect of *TricB* expression. We generated mice homozygous for inducible *Pkd2*-floxed allele with or without inducible conditional *TricB*-expressing allele. *TricB* transgene expression induced at the same time of *Pkd2*-cKO prevented mitochondrial morphological alternations at the pre-cystic and cystic stage in both PTs and DTs (Fig. 1, E and F; Fig. S1, A and B).

**Figure 1.**
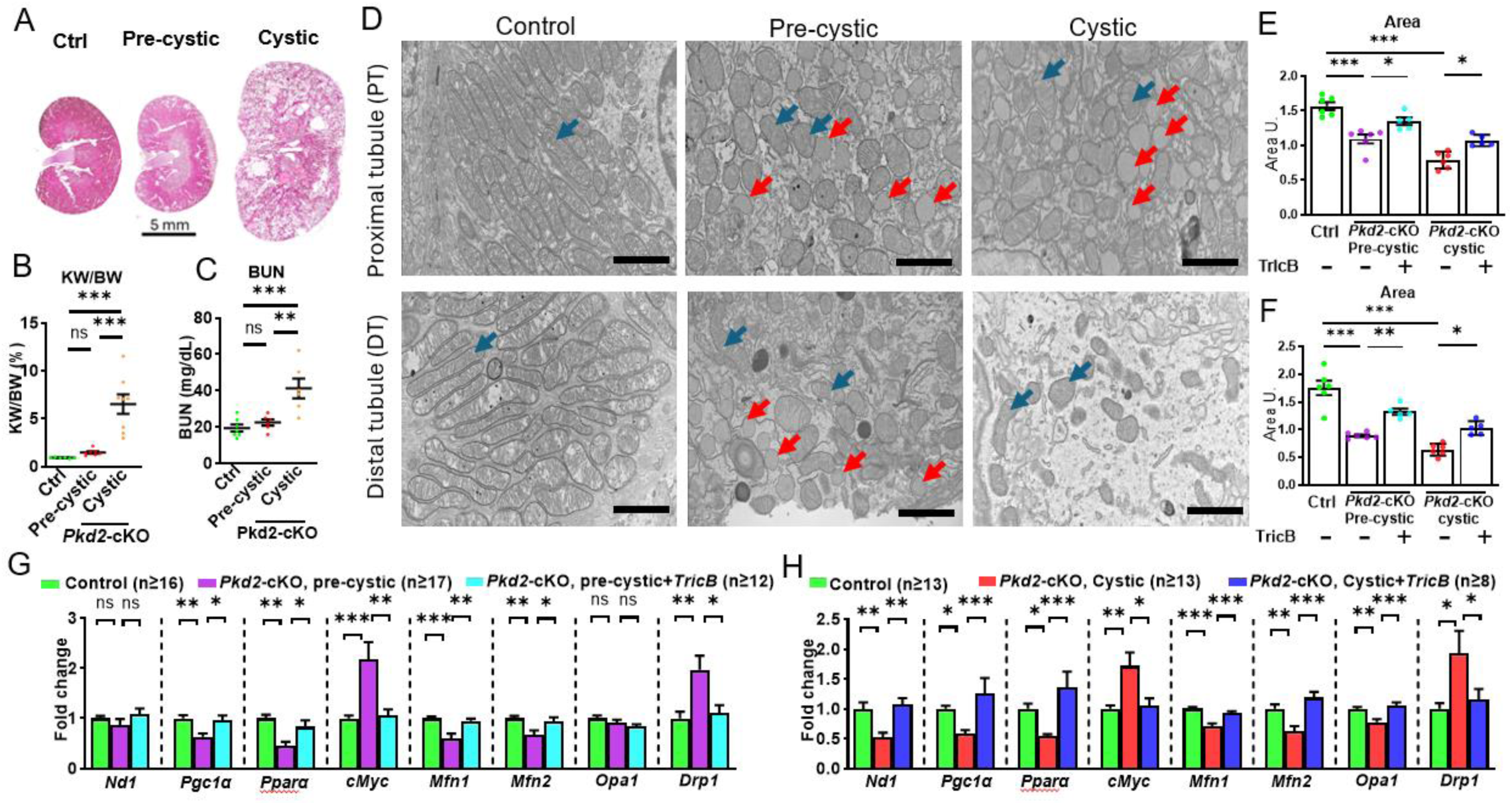
Kidney histology, mitochondrial morphological and gene expression changes in *Pkd2*-cKO mice at pre-cystic and cystic stages. (**A**-**C**) H&E-stained kidney sections (**A**), kidney weight/body weight ratio (**B**), and blood BUN levels (**C**) from pre-cystic (4 weeks post-induction) and cystic (12-16 weeks) kidneys of doxycycline-inducible kidney-specific *Pkd2*-cKO vs control. Scale bars for panel **A**, 5 mm. (**D**), Representative TEM images of proximal tubules (PTs, upper panel) and distal tubules (DTs, lower panel) from control vs pre-cystic and cystic *Pkd2*-cKO. Magnification at 10k. Blue and red arrows denote mitochondria and lysosomes, respectively. Note that the electron density of abnormal mitochondria is reduced. Electron density reflects heavy metals (used in TEM staining) binding to biological materials, lipids, proteins, DNA, etc. Scale bars, 2µm at 10k magnification. (**E, F**) Average mitochondrial area in control, pre-cystic and cystic *Pkd2*-cKO kidneys with or without *TricB* expression in PTs (**E**) and DTs (**F**). Each data point is the mean of >30 mitochondria (from multiple images) of one mouse. Each bar represents 5-7 mice per group. Data are presented as mean ± SEM. One-way ANOVA followed by Tukey’s multiple comparison test. (**G, H**) Quantitative real-time PCR analysis of genes in kidney tissues dissected from control and *Pkd2*-cKO ± *TricB* at the pre-cystic (**G**) and cystic stage (**H**). Data are mean ± SEM. P-values were calculated by one-way ANOVA followed by Tukey’s multiple comparison test. *, **, ***; p < 0.05, 0.005, 0.001, respectively. ns- not significant.

The mitochondrial biogenesis master regulator PGC1α is downregulated in *Pkd*-mutant mice,^12^ which would contribute to the mitochondrial biogenesis, structure, and function defects. ER Ca^2+^ release controls cytosolic [Ca^2+^] and regulates the expression of *Pgc1α* through calmodulin-dependent kinase II (CaMKII) and calcineurin (CaN).^13^ We examined whether the effect of *TricB* transgene might be through reversing *Pgc1α* expression. Compared to the controls, *Pgc1α* was reduced in cKO kidneys (Fig. 1, G and H). The number of mitochondria per cell analyzed by the mitochondrial/nuclear DNA ratio (ND1/HK2, labeled “*Nd1*”) was also significantly reduced in cKO kidneys vs controls. The peroxisome proliferator-activated receptor-α (*Pparα*) is a downstream gene target of PGC1α that controls multiple enzymes involved in fatty acid oxidation. As for *Pgc1α*, *Pparα* was significantly reduced in cKO kidneys vs controls. *cMyc* is an early gene that promotes cyst formation and is upregulated in ADPKD. We found *cMyc* was upregulated in cKO kidneys vs controls. Mitochondria undergo dynamic fusion and fission regulated by pro-fusion and pro-fission factors.^14^ The pro-fusion genes mitofusin-1 (*Mfn1*), mitofusin-2 (*Mfn2*), and optic atrophy 1 (*Opa1*) were downregulated in *Pkd2*-cKO pre-cystic and cystic kidneys. Conversely, the expression of pro-fission gene dynamin-related protein 1 (*Drp1*) was upregulated. Importantly, all these changes in mitochondrial biogenesis and dynamic regulatory genes were already significant at the pre-cystic stage (Fig. 1, G and H). *TricB* transgene expression reversed these changes in both pre-cystic and cystic *Pkd2*-cKO kidneys, restoring the levels to the control (Fig. 1, G and H). Overall, this gene signature supports mitochondrial structural and functional changes observed in *Pkd2*-cKO and mechanistically is related to ER Ca^2+^ homeostasis defects evidenced by the reversal by *TricB*.

### Deletion of Pkd2 alters mitochondria-associated ER membranes

Under normal conditions, the ER and mitochondria form close contacts as MAMs, which facilitate inter-organelle crosstalk. The global mitochondrial alterations observed in *Pkd2*-cKO kidneys likely reduce the frequency of these close-range contacts. However, MAMs are not entirely absent in cKO kidneys. We analyzed the structural features of MAMs in *Pkd2*-cKO versus control kidneys. Specifically, we focused on MAMs formed between the smooth ER and mitochondria with a separation distance of ≤30 nm, which are critical for direct Ca²⁺ transfer between the organelles without diffusion through the bulk cytosol.^15^ In addition to measuring the inter-organelle distance, we quantified the length of ER-mitochondria contacts (Fig. 2A). Representative images illustrating MAMs distances in proximal tubules (PTs) of cystic *Pkd2*-cKO kidneys versus controls, and the effect of *TricB* expression, are shown in Fig. 2B. Detailed analysis revealed a significant increase in MAMs distance and a decrease in contact length in both PTs and distal tubules (DTs) of pre-cystic and cystic kidneys compared to controls (Fig. 2, C–F). These findings indicate that although some MAMs contacts persist in cKO kidneys, their structural connectivity is compromised. Notably, the progression from pre-cystic to cystic stages showed minimal further changes in MAMs distance or length, suggesting that the pathology is substantially advanced prior to cyst formation. Importantly, *TricB* expression reversed the alterations in both MAMs distance and contact length in PTs and DTs of *Pkd2*-cKO kidneys. Previous studies, including our own, have confirmed that TricB is localized specifically to the ER.^6,16^

**Figure 2.**
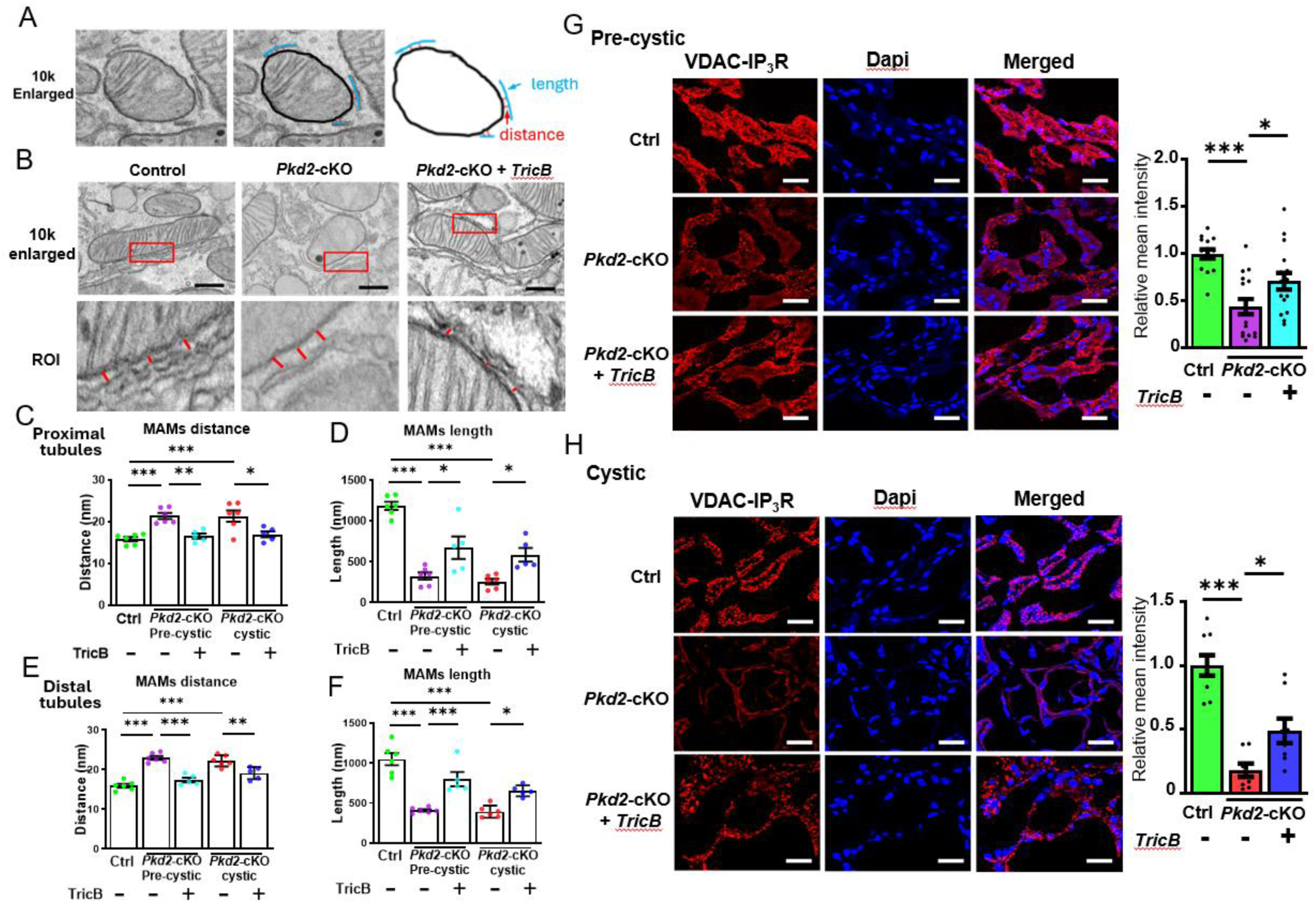
Alterations of MAMs structure in *Pkd2*-cKO mice and reversal by *TricB* expression. (**A**) Illustration of measurements of MAMs structure parameters in *Pkd2*-cKO. TEM images at enlarged 10k magnification. See text for description. (**B**) Representative TEM images depicting MAMs distance in control vs *Pkd2*-cKO cystic kidneys with or without *TricB* expression. Top, magnification at 10k. Scale bars, 0.5µm. Bottom, blown-up view of region of interest (ROI) marked in red box. (**C-F**) Mean MAMs distance and MAMs length in proximal tubules (**C** and **D**, respectively) and distal tubules (**E** and **F**, respectively) from control, pre-cystic, and cystic *Pkd2*-cKO mice with or without *TricB* expression. Note that the measurements focus on MAMs between smooth ER and mitochondria (separation distance ≤30 nm) ^9^. Each data point is the mean of >25 mitochondria (from multiple images) of one mouse. Each bar represents 5-7 mice per group. Data are mean ± SEM. P-values were calculated by one-way ANOVA followed by Tukey’s multiple comparison test. (**G**, **H**) Proximity ligation assays (PLA) probing the proximity between IP_3_R residing in ER and VDAC residing in OMM. Proximity <40 nm allows chemical reaction generating red fluorescent signals.^49^ Kidney sections were analyzed *in situ*. Nuclei were marked by Dapi staining (blue). Left panels are representative *in situ* PLA images in pre-cystic (**G**) and cystic kidneys (**H**). Right panels, bar graphs for PLA signals as relative mean intensity from 3 images per mouse from 5 mice in each group. Data are mean ± SEM. One-way ANOVA followed by Tukey’s multiple comparison test. Scale bars, 20µm.

The preceding TEM analysis focusing on structural alterations of existing MAM contacts does not account for the diminished ER-mitochondria crosstalk resulting from reduced mitochondrial number and size. To more comprehensively assess the impact of *Pkd* mutation on ER-mitochondria connectivity, we employed a proximity ligation assay (PLA). This technique quantifies the overall interaction between the ER and the outer mitochondrial membrane using the inositol trisphosphate receptor (IP_3_R) and voltage-dependent anion channel (VDAC) as respective probes. PLA offers additional advantages, including the elimination of operator bias inherent to TEM and the ability to evaluate all tubular segments beyond PTs and DTs. As shown, *Pkd2* deletion markedly reduced the IP_3_R-VDAC interaction in the pre-cystic as well cystic kidneys (Fig. 2, G and H). *TricB* expression partially reversed the decreased interaction in *Pkd2*-cKO kidneys. Overall, the results support the notion that disruption in ER-mitochondria connection occurs early in *Pkd2*-cKO and the role of ER Ca^2+^ homeostasis defects in the pathogenesis.

### Pkd2-cKO alters mitochondrial membrane potentials, decreases mitochondrial Ca^2+^ levels and mitochondrial respiration and reversal by correcting ER Ca^2+^ homeostasis by TricB expression

The mitochondrial membrane potential (ΔΨm) is generated by the activity of ETC complexes I and III, and IV, and an essential component in the process of energy storage during OXPHOS. ΔΨm together with the proton gradient (ΔpH) form the energy source for ATP synthesis by the ATP-synthetase (complex V). The levels of ΔΨm are normally kept relatively stable. A long-lasting change of ΔΨm’s reflects pathologies and serves as a signal for selective elimination of dysfunctional mitochondria. We measured mitochondrial membrane potentials in PTs isolated from *Pkd2*-cKO and control kidneys. Tetramethylrhodamine methyl ester (TMRM) is a cell-permeant potential-sensitive red fluorescent dye that selectively accumulates in metabolically active mitochondria.^17^ Selective partition to mitochondria was verified by colocalization with the mitochondrial marker MitoTracker green (MitoG) (Fig. 3A). ΔΨm in PTs of *Pkd2*-cKO kidneys was much higher (hyperpolarization) than in those of control kidneys. *TricB* expression in *Pkd2*-cKO normalized the ΔΨm. Notably, mitochondrial hyperpolarization has also been observed by others in *Pkd*-mutant cell lines.^5,18^ To explain why ΔΨm increases despite reduced mitochondrial activity, it’s important to consider the underlying bioenergetic balance. ΔΨm reflects the equilibrium between its generation by ETC complexes I, III, and IV, and its dissipation through ATP synthesis by complex V. Although the activity of all ETC complexes is diminished in *Pkd2*-mutant mice, a disproportionately greater reduction in complex V activity—relative to complexes I, III, and IV—may lead to reduced proton flux through ATP synthase. This imbalance could result in proton accumulation across the inner mitochondrial membrane and mitochondrial hyperpolarization.

**Figure 3.**
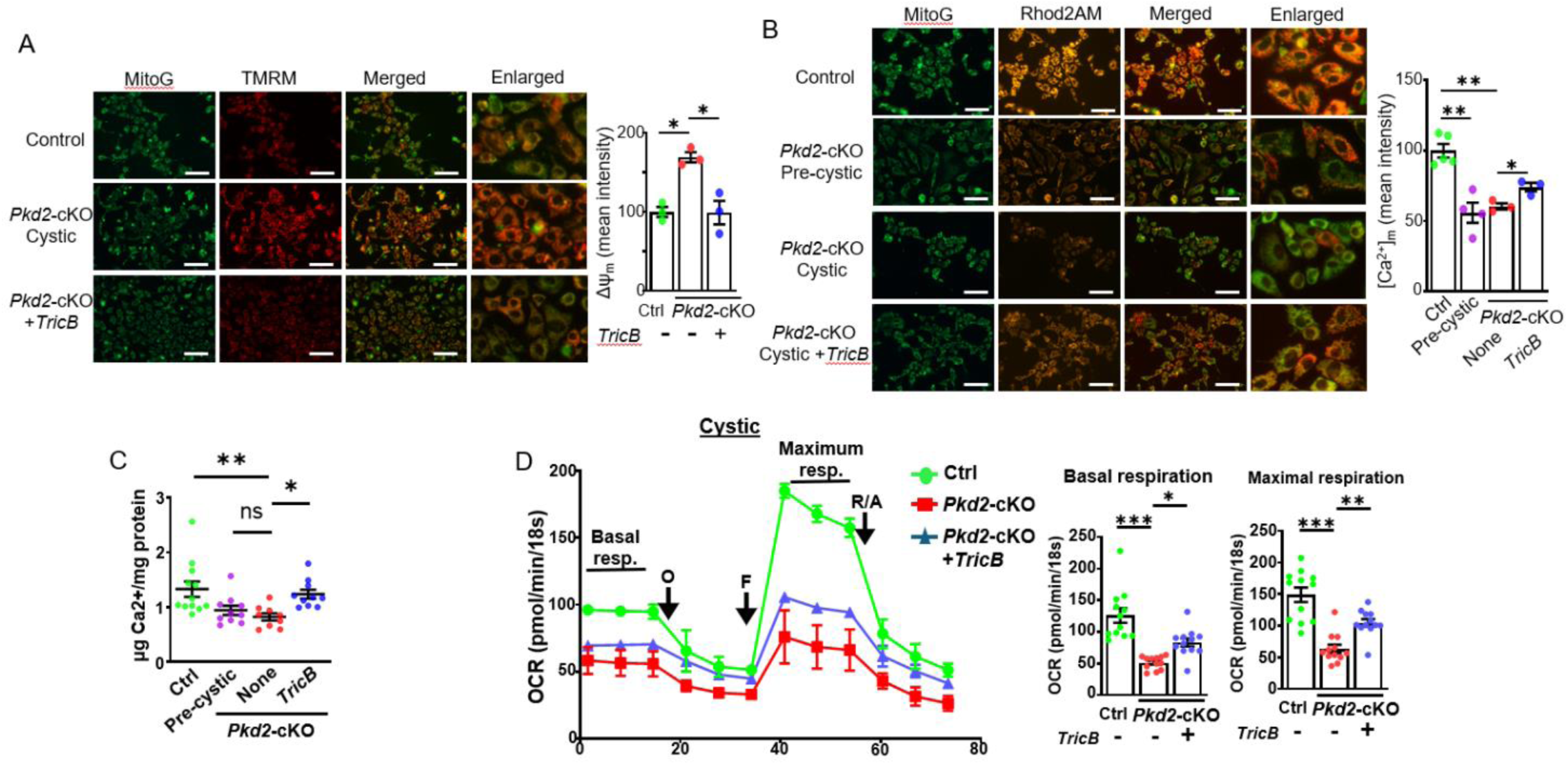
Alterations of mitochondrial membrane potential, calcium concentration, calcium content, and oxygen consumption rate in proximal tubules isolated from *Pkd2*-cKO mice and reversal by *TricB*. (**A**) Mitochondrial membrane potentials (ΔΨm) were measured in PTs from control, *Pkd2*-cKO cystic kidneys with and without *TricB*. PTs from control and non-cystic regions of *Pkd2*-cKO kidneys with or without TricB were freshly isolated and incubated in HBSS buffer containing the potential-sensitive dye TMRM and the mitochondrial marker MitoG. Left panels are representative overlaid phase contrast (DIC) and TMRM fluorescence (Cy3 channel) images. The far-right column shows enlarged images. Scale bars, 200µm. Right panel is bar graph of ΔΨm from corresponding groups. Control (Ctrl) is set as 100%. Mean ± SEM of 3 mice. Data from each mouse are averaged from at least 10 separate measurements. (**B**) Mitochondrial Ca^2+^ concentration ([Ca^2+^]) in PTs from control, *Pkd2*-cKO cystic kidneys with and without *TricB* were measured. PTs were incubated with Rhod2AM, which enters cytosol then cellular organelles in the acetoxymethyl ester form and is retained after esterase cleavage. Preferential retention of Rhod2 in mitochondria is achieved by selected permeabilization of the cell and ER membrane by saponin treatment followed by washing to remove dyes in the latter compartments.^17^ Throughout the preparation, isolated PTs were bathed in a solution in which [Ca^2+^] was buffered at 100 nM. Left panels were representative overlaid DIC and Rhod2 fluorescence images. Scale bars, 200µm. The right panel is the bar graph of [Ca^2+^] from corresponding groups. Mean ± SEM of 3-6 mice followed by unpaired t-test with Welch’s correction. Data from each mouse are averaged from at least 10 separate measurements. (**C**) Mean ± SEM of calcium contents of mitochondria isolated from control, *Pkd2*-cKO pre-cystic, and cystic kidneys with or without *TricB* expression. One-way ANOVA followed by Tukey’s multiple comparison test employed. (**D**) OCR of PTs isolated from control and *Pkd2*-cKO cystic kidneys with or without TricB. O, F, R, and A are mitochondrial ETC inhibitors oligomycin, FCCP (carbonyl cyanide 4-[trifluoromethoxy]phenylhydrazone), rotenone, and antimycin A, respectively. The left panel shows representative line tracings of one experiment. Data points in each group represent mean ± SD of multiple wells from a single mice per group with each well containing 9 -10 freshly dissected PTs. OCR values were normalized to 18S rRNA measured by rt-PCR. Bar graphs on the right represent mean ± SEM of >10 single wells with at least 7 mice per group. Basal respiration is OCR before addition of ATP synthetase inhibitor oligomycin. Maximal respiration is OCR after addition of mitochondrial OXPHOS uncoupler FCCP. P-values were calculated by one-way ANOVA followed by Tukey’s multiple comparison test. *, **, ***; p < 0.05, 0.005, 0.001, respectively. ns- not significant.

Mitochondria are also important regulators of cellular Ca^2+^ homeostasis by sequestering and releasing Ca^2+^.^19^ Mitochondrial [Ca^2+^] also regulates mitochondrial metabolism, ATP production, and cell death.^19^ Mitochondrial [Ca^2+^] was measured using a calcium-sensitive fluorescent dye rhodamine-2-acetoxymethyl ester (Rhod2 AM).^20^ As shown, mitochondrial [Ca^2+^] was reduced in both pre-cystic and cystic *Pkd2*-cKO PTs relative to the control (Fig. 3B). *TricB* expression partially reversed the decreases in mitochondrial [Ca^2+^] in *Pkd2*-cKO PTs. To further support that [Ca^2+^] measurement by Rhod2 reflects mitochondrial [Ca^2+^], we isolated mitochondria from control and *Pkd2*-cKO kidneys to measure Ca^2+^ content. Mitochondrial Ca^2+^ content was significantly reduced in the pre-cystic and cystic *Pkd2*-cKO kidneys compared to control (Fig. 3C). Mitochondria from *Pkd2*-cKO with transgenic *TricB* expression showed partially restored mitochondrial Ca^2+^ content vs those in *Pkd2*-cKO without *TricB* expression. The mitochondrial respiratory function was studied by measuring oxygen consumption rate (OCR) (Fig. 3D). OCR assay revealed a decrease in OXPHOS activity in PTs isolated from *Pkd2*-cKO cystic kidneys compared to the control (Fig. 3D). Both normalized basal respiration and maximal respiration were significantly reduced in *Pkd2*-cKO compared to controls. Transgenic *TricB* expression partially restored OCR in the PTs of *Pkd2*-cKO mice evidenced by improvements in both basal and maximal respiration. For pre-cystic kidneys, OXPHOS activity measured at the basal and in the presence of mitochondrial inhibitors revealed a declining trend but did not reach statistical significance (Fig. S2). As discussed below, isolated tubule studies under in vitro setting may not completely reflect the pathology in native kidneys. These functional studies nonetheless are consistent with the above genetic and structural analysis of mitochondrial alterations in *Pkd*-mutants.

### Pkd1-cKO share similar mitochondrial structural and functional changes with Pkd2-cKO and correction by TricB expression

Mutations of *PKD1* and *PKD2* in humans phenocopy each other barring a difference in the age at the onset of disease in affected individuals.^1^ While the precise mechanisms underlying the phenocopy remain elusive, it begs the question whether *Pkd1*-cKO shares similar mitochondrial pathology and responsiveness to *TricB* expression as *Pkd2*-cKO. As shown, cystic *Pkd1*-cKO kidneys showed markedly enlarged cystic kidneys while co-expressing *TricB* transgene ameliorated cyst formation with decreased kidney size (Fig. 4, A and B). Cyst index and BUN levels were elevated in *Pkd1*-cKO and partially reversed by *TricB* expression (Fig. 4, C and D). Thus, as is for *Pkd2*-cKO mice,^6^ *TricB* expression suppresses cystogenesis in *Pkd1*-cKO. Similar to *Pkd2*-cKO kidneys, mitochondria in *Pkd1*-cKO kidneys were fewer, smaller in size and rounded with decreased cristae density in both pre-cystic and cystic mice compared to controls (Fig. S3, A and B). These findings were supported by quantitative analysis of mitochondrial morphology and mitochondrial area in TEM images of *Pkd1*-cKO kidneys. *TricB* expression in *Pkd1*-cKO reversed mitochondrial alterations in pre-cystic as well as cystic kidneys. As is for *Pkd2*-cKO, the number of mitochondria per cell as reflected by mitochondrial/nuclear DNA ratio, master regulators for mitochondria biogenesis and function, and genes regulating mitochondrial dynamics were altered in *Pkd1*-cKO kidneys in pre-cystic and cystic stages and reversed by *TricB* expression (Fig. 4, E and F).

**Figure 4.**
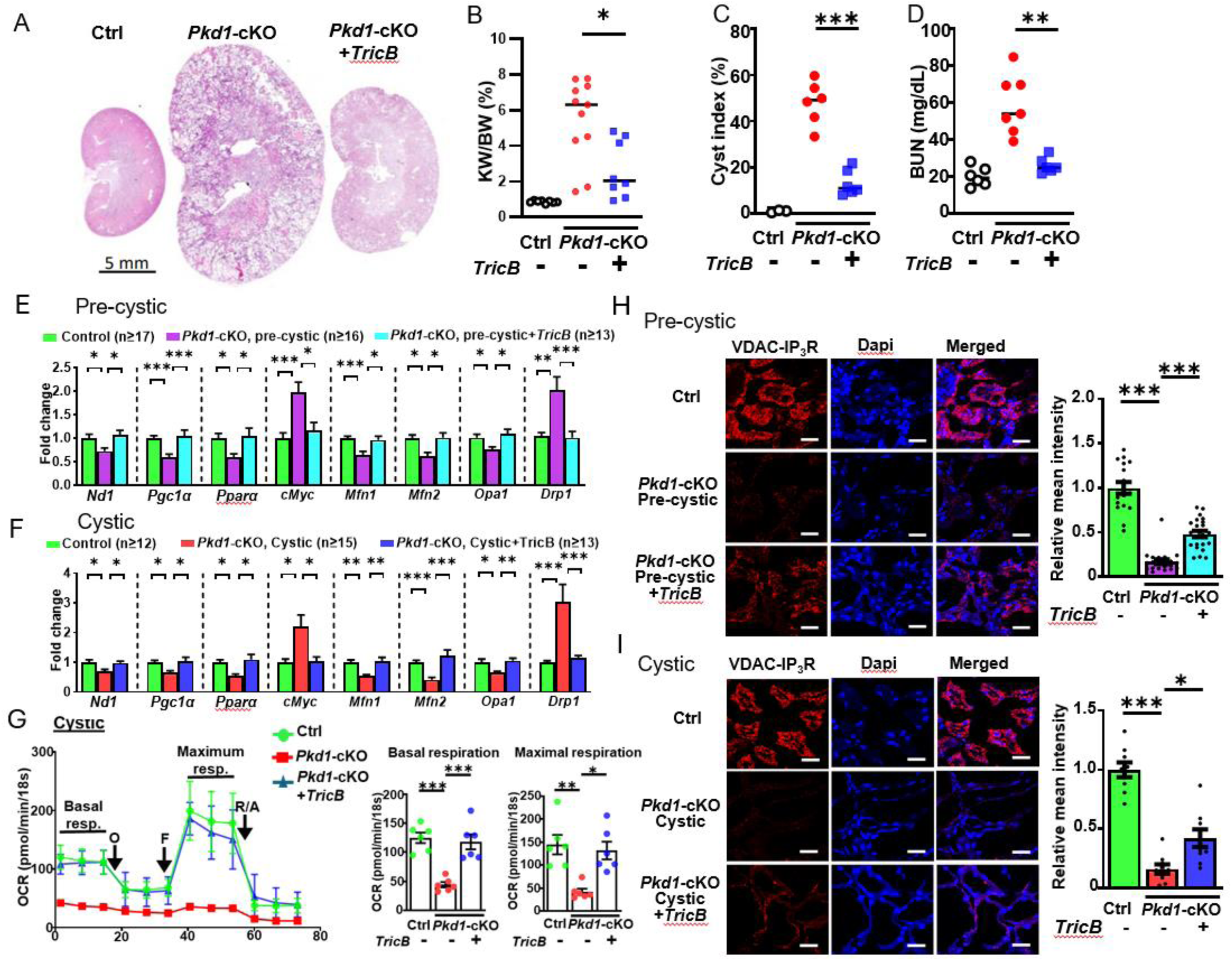
*TricB* expression reverses cystic phenotypes and mitochondrial alterations in *Pkd1*-cKO mice. (**A**) Representative H&E-stained kidney sections of control, *Pkd1*-cKO ± *TricB* mice. Scale bar is 5 mm. Mice carrying doxycycline inducible kidney-specific *Pkd1*-floxed allele (*Pkd1^f/f^*;*Pax8-LC1*) with or without the allele for inducible kidney-specific expression of *TricB* transgene (*cTg-TricB*) were studied. (**B**-**D**) Mean ± SEM of kidney weight/body weight ratio (**B**), cyst index (**C**), and blood BUN levels (**D**) of control, *Pkd1*-cKO ± *TricB* mice. (**E**, F) Fold change of targeted gene expressions analyzed by qRT-PCR in *Pkd1*-cKO ± *TricB* at the pre-cystic (**E**) and cystic stage (**F**). See Fig. **1, G** **&H** for descriptive details. (**H**, **I**) PLA analysis of ER-mitochondria connection in *Pkd1*-cKO ± *TricB*. Left panels are representative *in situ* PLA images in pre-cystic (**H**) and cystic kidneys (**I**). Right panels, bar graphs for PLA signals as relative mean intensity from 3 images per mouse from 5 mice in each group. Data are mean ± SEM. Assessed by one-way ANOVA with Tukey’s multiple comparison test. *, **, ***; p < 0.05, 0.005, 0.001, respectively. ns- not significant.

Further examination of mitochondrial function showed a significant reduction of OCR in PTs isolated from the cystic stage of *Pkd1*-cKO kidneys (Fig. 4G). *TricB* expression restored OCR to the level of control kidneys. In pre-cystic PTs, OCR showed a trend to decrease but was not significantly different from controls (Fig. S4). As was in *Pkd2*-cKO, the downregulation of *Pgc1α* and *Mfn2* in *Pkd1*-cKO predicts that MAMs structure would be altered. TEM revealed an increased MAMs distance and reduced MAMs length in pre-cystic and cystic *Pkd1*-cKO and reversal by *TricB* expression (Fig. S3, B and C). The ER-mitochondria connectivity measured by PLA also showed reduced interactions in *Pkd1*-cKO pre-cystic and cystic kidneys and reversal by *TricB* expression (Fig. 4, H and I). These results highlight those disruptions in Ca^2+^-dependent ER-mitochondria connection leading to mitochondrial dysfunction is an early denominator for cystogenesis in both *Pkd1*- and *Pkd2*-cKO.

### Enhancing MAM function by Sigma-1 receptor agonist reverses mitochondrial dysfunction and cystogenesis in Pkd1-cKO

We have shown ER-mitochondria connection is disrupted early in the pre-cystic stages and that mitochondrial structural and functional changes and cystogenesis in *Pkd1*- and *Pkd2*-cKO are reversed by correcting ER Ca^2+^ release defect by *TricB*. Sigma-1 (σ1) receptor (S1R) is a membrane-spanning Ca^2+^-sensitive ER chaperone protein predominantly localized to MAMs.^21,22^ Pre-084 is a highly selective S1R activator that enhances MAMs function to improve mitochondrial function and neurodegenerative disease phenotypes.^23^ We examined whether enhancing MAMs function by pre-084 can ameliorate mitochondrial dysfunction and cystogenesis in *Pkd*-mutant mice. We focused on *Pkd1*-cKO mice as they develop cysts more aggressively. As shown, *Pkd1*-cKO mice receiving pre-084 had smaller kidneys with few cysts compared to saline injected mice (Fig. 5, A-C). BUN levels were significantly lower in pre-084-treated than saline-treated mice (Fig. 5D). TEM analysis revealed improved mitochondrial and MAMs structures in pre-084-treated kidneys vs untreated kidneys (Fig. S5, A-D).

**Figure 5.**
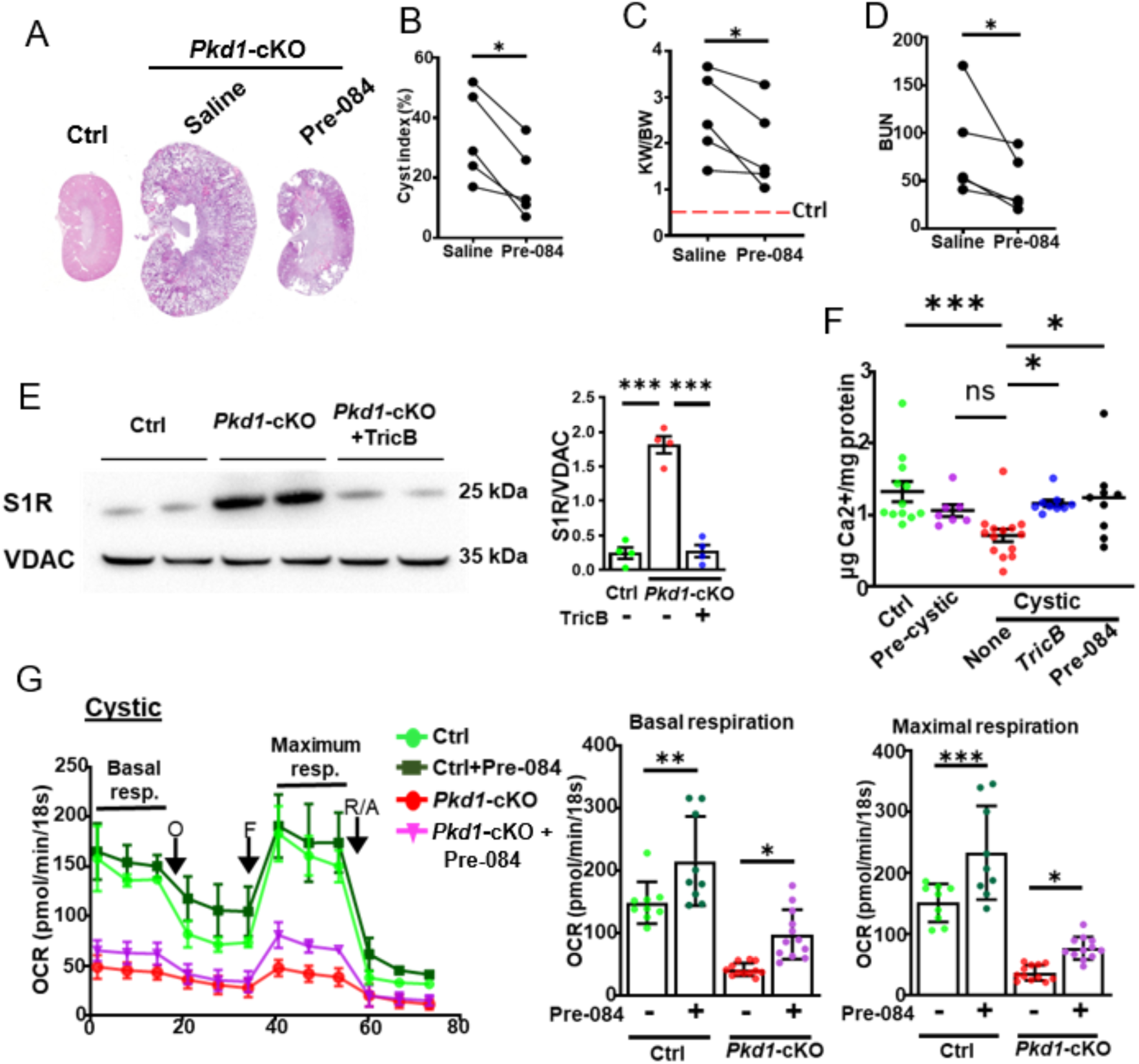
Pre-084 reverses cystic phenotypes, mitochondrial morphological and functional alterations in *Pkd1*-cKO mice. (**A**) Representative H&E-stained kidney sections of control (Ctrl), and *Pkd1*-cKO treated with saline (vehicle) or pre-084. *Pkd1*-cKO mice received daily intraperitoneal injection of pre-084 (1 mg/kg daily) or saline at the completion of doxycycline induction. Kidneys were harvested at 12 weeks after induction. (**B**-**D**) Cyst index (**B**), KW/BW ratio (**C**), and blood BUN levels (**D**) of control and *Pkd1*-cKO mice treated with saline or pre-084. Each data point is an average of 2-3 mice from saline or pre-084-treated group in each separate experiment. A total of 5 separate experiments with *Pkd1*-cKO mice treated with saline or pre-084 were conducted. Paired student *t*-test. *, p< 0.02. (**E**) Representative western blot of sigma-1 receptor (S1R) protein in MAMs fraction isolated from control (Ctrl), *Pkd1*-cKO cystic kidney ± *TricB*. VDAC localized to the outer mitochondria membrane of MAMs is used as protein loading control. Right panel-graph represents mean S1R protein expression in *Pkd1*-cKO mice with or without TricB in comparison to control, n=4. One-way ANOVA followed by Tukey’s multiple comparison test. ***; p < 0.001. (**F**) Ca^2+^ content in mitochondria isolated from control, *Pkd1*-cKO pre-cystic, and *Pkd1*-cKO cystic kidneys with or without *TricB* expression, and *Pkd1*-cKO treated with pre-084. Pre-084 was administered to *Pkd1*-cKO mice soon after doxycycline induction till sacrifice. P-values were calculated by one-way ANOVA followed by Sidak’s multiple comparison test. *, **, ***; p < 0.05, 0.005, 0.001, respectively. ns- not significant. (**G**) OCR measurement in control and *Pkd1*-cKO PTs ± pre-084. Freshly isolated PTs were incubated with saline or pre-084 for 2 hrs before OCR measurement. The left panel shows a representative line plot from one experiment. Data points in each of 4 groups represent mean ± SD of 3 wells with each well containing 9-10 freshly dissected PTs. OCR values are normalized to 18S rRNA measured by rt-PCR. Bar graphs on the right represent mean ± SEM of 3 individual experiments. One-way ANOVA followed by Sidak’s multiple comparison test. Please note that the modest effect of pre-084 on *Pkd1*-cKO tubules (∼20% increases) may be a result of short-term incubation *in vitro*.

ER chaperon proteins may be upregulated in response to stress.^21,24^ Indeed, S1R was markedly upregulated in *Pkd1*-cKO kidneys, and *TricB* expression reversed *Pkd1*-cKO-induced upregulation (Fig. 5E). S1R normally forms inactive complexes with the binding immunoglobulin protein (BiP), another chaperone commonly upregulated during ER stress. S1R agonist dissociates the complexes allowing S1R to bind IP_3_R and promote Ca^2+^ release.^21,22,25^ Thus, simple upregulation of S1R is not sufficient to confer increased function and activation by an agonist is required. To support that pre-084 enhances mitochondrial function, we found that acute treatment of isolated PTs from *Pkd1*-cKO with S1R agonist pre-084 augmented the basal and maximal mitochondrial respiration (Fig. 5G). While control PTs have higher OCR, acute treatment with pre-084 further increased the activities. To support that pre-084 acts on MAMs to promote ER-mitochondria Ca^2+^ transfer, we administered pre-084 to *Pkd1*-cKO mice and determined Ca^2+^ content in isolated mitochondria. Pre-084 treatment partially reversed the decrease in mitochondrial Ca^2+^ content in *Pkd1*-cKO (Fig. 5F). For comparison, *TricB* expression also partially reversed the decrease in mitochondrial Ca^2+^ content in *Pkd1*-cKO.

### ADPKD mice have altered epigenetic landscape marked by chromatin histone acetylation

Widespread transcriptional dysregulation is a feature of ADPKD. Mitochondrial metabolic disturbances in ADPKD alter the production of acetyl-coenzyme A (acetyl-CoA), an essential cofactor for epigenetic modification histone acetylation.^26^ We asked whether mitochondrial dysfunction resulting from defects in ER Ca^2+^ homeostasis in *Pkd*-cKO may alter epigenetic landscape contributing to the transcriptome dysregulation. Nuclear lysates from *Pkd1*- and *Pkd2*-cKO kidneys had increased levels of histone H3K27 acetylation (H3K27ac), a marker for active enhancers (Fig. 6A). The increases were evident in both pre-cystic and cystic kidneys and reversed in *Pkd*-cKO by *TricB* expression or treatment with pre-084. *cMyc* is an oncogene upregulated in ADPKD that plays a key role in its pathogenesis.^27^ Lakhia *et al.,* recently reported that histone acetylation of *cMyc* enhancers is important in activating the gene.^26^ We examined several H3K27ac marked-*cMyc* enhancer regions (E5, E19 and E28) and found *TricB* transgene expression or pre-084 treatment reversed the increases in these marks in both *Pkd1*- and *Pkd2*-cKO (Fig. 6, B-D). This provides compelling support for the notion that ER-mitochondria disruption contributes to ADPKD pathogenesis by driving epigenetic rewiring and *cMyc*-regulated cell events.

**Figure 6.**
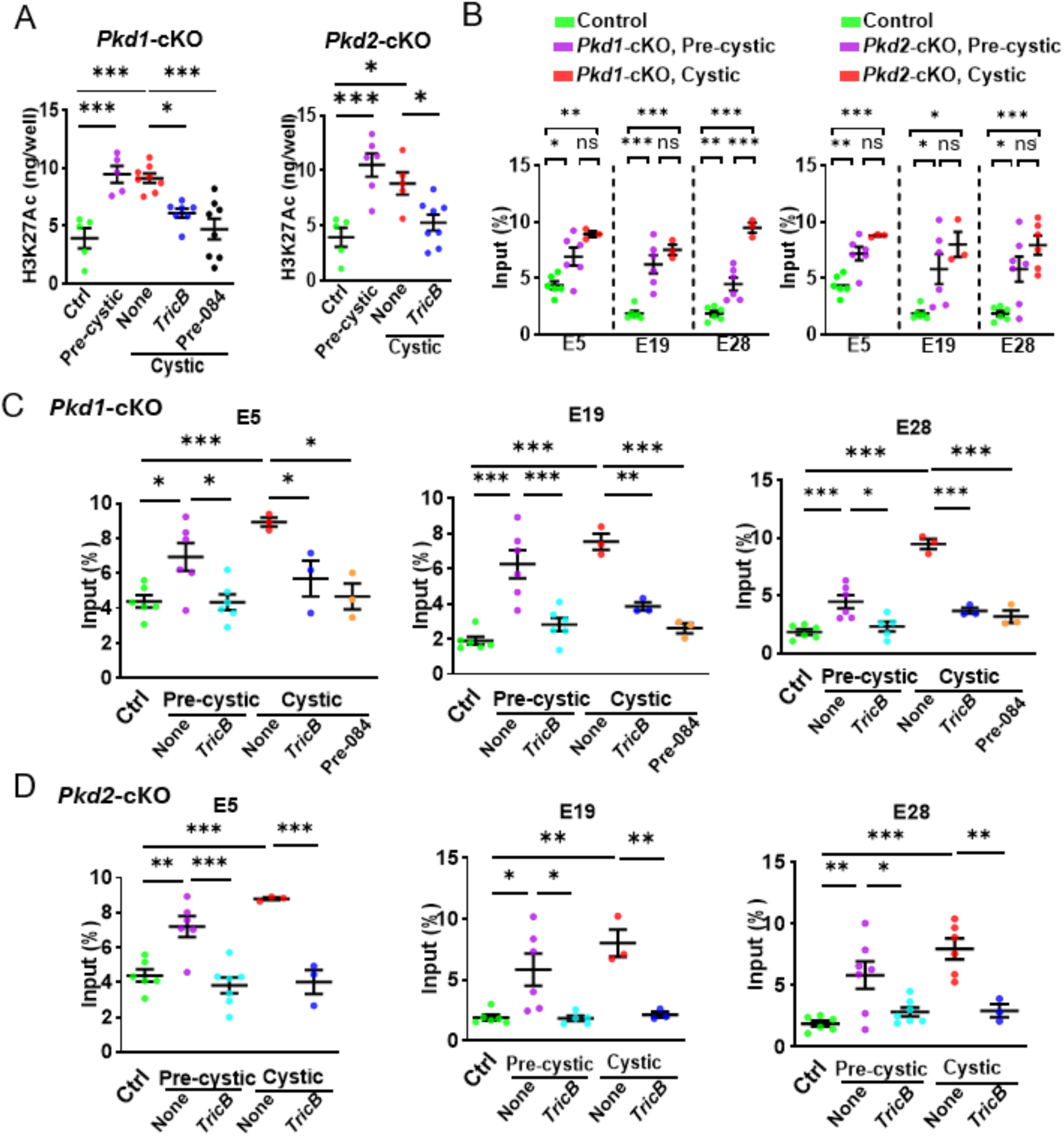
Epigenetic landscape of *cMyc* in *Pkd1*-cKO and *Pkd2*-cKO revealed by chromatin histone H3K27 acetylation. (**A)** Total H3K27 acetylated histone level in kidney lysates were measured by ELISA from control kidney, *Pkd*-cKO pre-cystic kidney, *Pkd*-cKO cystic kidney ± *TricB* or treated with pre-084. Data are mean ± SEM. One-way ANOVA followed by Sidak’s multiple comparison test. (**B**-**D**) ChIP-qPCR analysis of histone H3K27 acetylation around *cMyc* genome locus in *Pkd*-cKO and effects of TricB and pre-084. ChIP-qPCR assay of histone H3K27 acetylation marked *cMyc* enhancers E5, E19 and E28 (as described in Lakhia et al.)^24^ in *Pkd1*-cKO (**B**, **C**) and *Pkd2*-cKO (**B**, **D**) with or without TricB expression or pre-084 treatment compared to control. Data are presented as mean ± SEM from 3 mice per group. P-values were calculated by one-way ANOVA followed by Sidak’s multiple comparison test. *, **, ***; p < 0.05, 0.005, 0.001, respectively. ns- not significant.

Recently, Mi *et al.,* reported that cyclin-dependent kinase-7 (CDK7) is upregulated in *Pkd*-mutant mice and required for the assembly of super-enhancer complexes for transcriptional activation of genes for metabolic reprogramming.^10^ Indeed, *Cdk7* gene expression was increased in the pre-cystic and cystic *Pkd1*- and *Pkd2*-cKO kidneys (Fig. 7, A and B). Importantly, the upregulation was reversed by *TricB* expression or treatment with pre-084 (Fig. 7, A and B). The annotated integrative genomics viewer shows numerous H3K27ac marks on genomic loci around the *Cdk7* gene (Fig. 7C, top) (http://genome.ucsc.edu). We selected thirteen H3K27ac marks (C1-C13) around *Cdk7* and measured their enrichment by chromatin-immunoprecipitation (ChIP)-qPCR. Fixed chromatin samples were immunoprecipitated by anti-H3K27ac antibodies. Eluted DNA was analyzed by qPCR analysis. Among the thirteen H3K27ac marks, eight loci were increased in pre-cystic and/or cystic kidneys of *Pkd1*-cKO, and five loci in *Pkd2*-cKO (Fig. 6C). This suggests a prominent role of CDK7 as a driver for metabolic reprogramming in *Pkd1*- and *Pkd2*-cKO mouse models.

**Figure 7.**
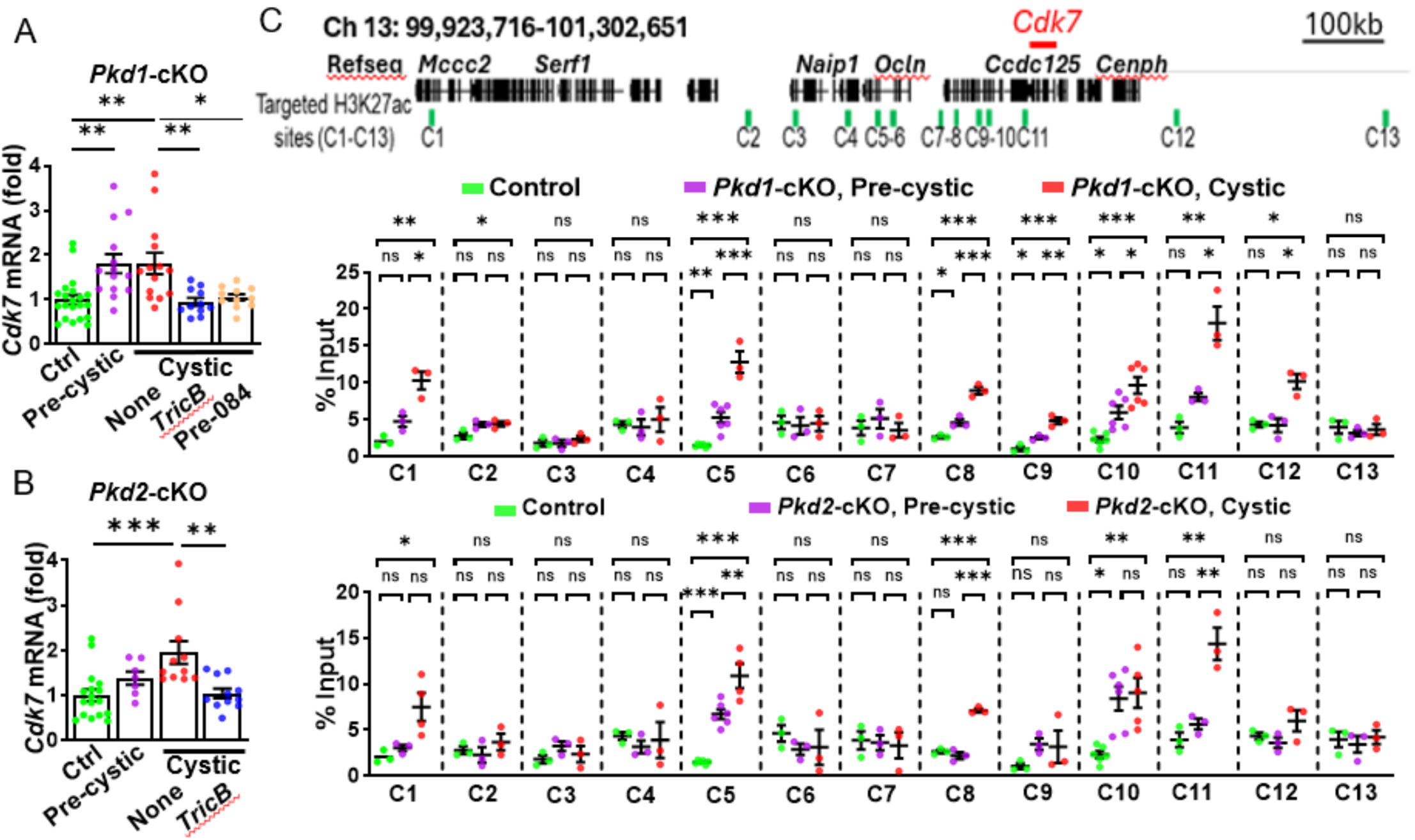
H3K27 histone acetylation mediates *Cdk7* upregulation and reversal of acetylation by *TricB* transgene or MAMs activator pre-084. (**A, B**) Quantitative real-time PCR analysis of *Cdk7* gene expression in control, *Pkd1*- and *Pkd2*-cKO pre-cystic kidneys and cystic kidneys ± *TricB* or treated with pre-084. Data are mean ± SEM. P-values were calculated by one-way ANOVA followed by Sidak’s multiple comparison test. (**C**) Top: genomic location of H3K27ac marks in Integrative Genomics viewer V2.18.4. Thirteen H3K27ac marks (C1 - C13) surrounding *Cdk7* gene selected for analysis. Bottom: ChIP-qPCR data illustrating the percentage input of targeted H3K27ac-marked genomic regions (C1-C13) in the kidneys of *Pkd1*-cKO (upper panel) and *Pkd2*-cKO (lower panel) at both pre-cystic and cystic stages, compared to control mice. Data are presented as mean ± SEM from 3-5 mice per group. P-values were calculated by one-way ANOVA followed by Tukey’s multiple comparison test. *, **, ***; p < 0.05, 0.005, 0.001, respectively. ns- not significant.

### Enhancer histone acetylation resulting from defects in ER-mitochondria connection mediates transcriptional upregulation of Cdk7

To investigate whether the open chromatin regions surrounding *Cdk7* are enhancers that drive the gene expression, we performed chromatin conformation capture (3C) assays to evaluate the interaction with the promoter (Fig. S6). Among the 5 putative enhancer regions that were identified in both *Pkd1*- and *Pkd2*-cKO, we selected C1, C5, and C10 for analysis. These regions show strong prediction as enhancers based on alignment with “K27ac kidney 16”, “ATAC kidney E15”, and “ENCODE cCREs” database (Fig. S6; Fig. 8A). Formation of a loop structure between enhancer and promoter permits the amplification of a PCR product despite a long linear distance separation (Fig. S6). As shown, the relative crosslinking/ligation frequency between the restriction fragment 11 in the C5 region (or the fragment 4 in the C10 region) and the anchor fragment marking the *Cdk7* promoter was significantly higher compared to other 3C restriction fragments in control kidneys (Fig. S6). The results indicate that the C5 and C10 are enhancer regions interacting with *Cdk7* promoter and playing a direct role in the transcriptional activation of *Cdk7* gene expression. For comparison, the C1 chromatin region did not show interaction with the *Cdk7* promoter; it may be an enhancer for other ADPKD upregulated genes. To support their roles in *Pkd*-mutant mice, we found the physical interaction between C5 and C10 and the *Cdk7* promoter was markedly increased in *Pkd1*-cKO cystic kidneys vs controls (Fig. 8A), supporting the notion that the H3K27ac-mediated epigenetic mechanism is important in the upregulation of *Cdk7* gene. To support the hypothesis that ER-mitochondria disconnection contributes to *Cdk7* upregulation, *TricB* transgene expression or the MAMs activator pre-084 reversed the increases in histone acetylation at C5 and C10 enhancer regions in *Pkd1*- and *Pkd2*-cystic kidneys (Fig. 8, B and C). The role of the C5 and C10 genomic regions as enhancers for *Cdk7* is further supported by the results that deletion of these regions in cultured mouse distal convoluted tubule (MDCT) cells resulted in reduced gene expression compared to the controls (Fig. 8D).

**Figure 8.**
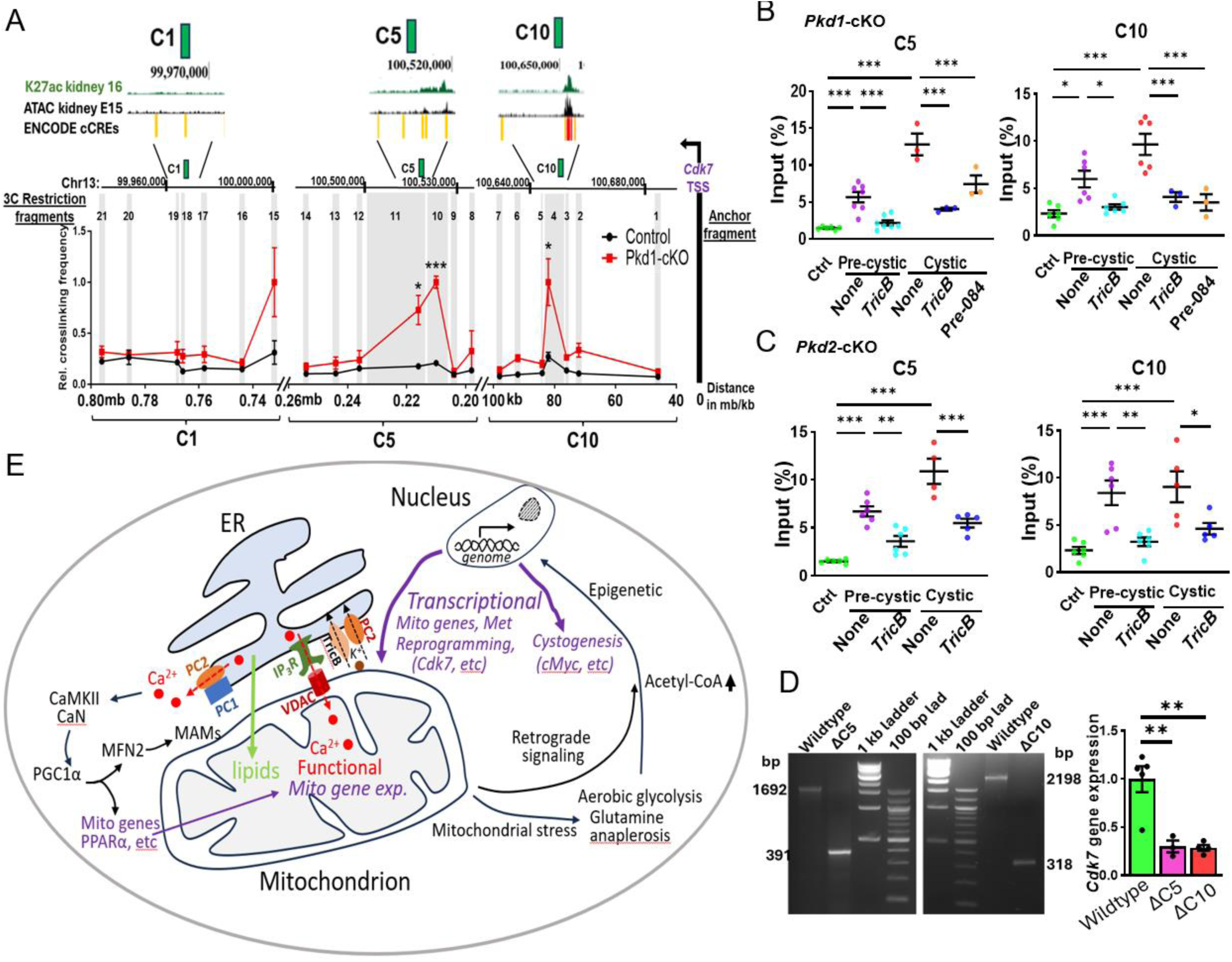
Confirmation of hyperactive enhancers of *Cdk7* in *Pkd*-mutant mice by chromosome conformation capture and validation by enhancer deletion analysis. (**A**) Top: enlarged illustration of H3K27ac marks depicted in the UCSC genome browser aligned with database of “K27ac kidney 16”, “ATAC kidney E15”, and “ENCODE cCREs”. K27ac kidney 16: kidney acetylated histone K27; ATAC kidney E15: “assay for transposase-accessible chromatin” kidney E15; ENCODE cCREs: ENCODE putative cis-regulatory elements. The C1 region is ∼0.75 mb whereas C5 and C10 regions are ∼100-200 kb away from the promoter. Bottom: relative crosslinking frequencies between targeted restriction fragments (light vertical gray bars) and the anchor fragment (black vertical bar to the right) tagging *Cdk7* promoter in control and *Pkd1*-cKO cystic kidneys. Ligation frequencies between restriction fragments were measured by qPCR per chromosome conformation capture assays (see Fig. S6). Total 21 restriction sites (producing 21 restriction fragments) with 7 each surrounding C1 (labelled as 21-15), C5 (14-8) or C10 (7-1) genomic regions were selected. Unpaired student *t*-test. (**B**, **C**) ChIP-qPCR assays targeting the C5 and C10 regions enriched with H3K27ac marks showed upregulation in *Pkd1*-cKO (**B**) and *Pkd2*-cKO (**C**) mice and reversed by *TricB* or pre-084 treatment. (**D**) Deletion of locus surrounding C5 (ΔC5) and C10 (ΔC10) enhancer regions in MDCT cells (gel image) leads to decrease in *Cdk7* gene expression compared to control (ctrl) (bar graph). Data are presented as mean ± SEM from 3-5 mice per group. P-values were calculated by one-way ANOVA followed by Tukey’s multiple comparison test. *, **, ***; p < 0.05, 0.005, 0.001, respectively. ns- not significant. (**E**) A working model illustrating the role of ER-mitochondria disconnection in pathogenesis of ADPKD. Loss of PCs-regulated ER Ca^2+^ release decreases mitochondrial Ca^2+^ concentration ([Ca^2+^]), leading to reduced TCA enzyme activities (“functional”). Decreases in cytosol [Ca^2+^], through CaMKII and CaN, downregulates the mitochondrial biogenesis master regulator PGC1α, which leads to downregulation of MFN2 and PPARα. Reduced activity of master regulators PGC1α and PPARα leads to downregulation of gene expression of TCA enzymes (“transcriptional”) and other mitochondrial biogenesis and dynamics-regulating genes. Mitochondrial membrane biogenesis depends on the transfer of lipids synthesized in the ER.^50^ Disruption in ER-mitochondria connection will further compromise mitochondrial membrane biogenesis. These functional and transcriptional alterations initiate mitochondrial stress and metabolic disruption, which are relayed to the nucleus through mitochondrial retrograde signaling. In response, mitochondria-driven epigenetic reprogramming activates a broad transcriptional program that orchestrates the diverse cellular processes essential for cystogenesis. See text for abbreviation and details.

## Discussion

PC1 and PC2 are present in multiple subcellular compartments. The precise function of PC1 and PC2 therein and how loss of function leads to cyst formation remain incompletely understood. We have previously shown that ER-localized PC2 contributes to anti-cystogenesis.^6^ IP_3_R-mediated ER Ca^2+^ release generates an ER lumen-negative potential which counters Ca^2+^ release. TricB is an ER-resident K^+^-permeable channel that mediates ER luminal K^+^ influx in exchange for Ca^2+^ efflux. Like TricB, PC2 is a non-selective cation channel more permeable to K^+^ than Ca^2+^.^7^ We showed that recombinant TricB complements defects in Ca^2+^ release in PC2-null cells, and expression of *TricB* transgene in *Pkd2*-cKO ameliorates cyst formation.^6^ Here, we further show that *TricB* transgene mitigates cyst formation in *Pkd1*-cKO.

Mitochondrial dysfunction and metabolic reprogramming occur in ADPKD and contribute to disease pathogenesis.^3–5,18^ How these alterations develop in ADPKD is unclear. The number of mitochondria in a renal tubular cell varies from hundreds to several thousands. Altogether they occupy ∼40% of the intracellular volume.^28^ Mitochondrial surface forms close contact with the ER membrane called mitochondrial associated membranes (MAMs). Extending our previous report that polycystins in the ER are important in anti-cystogenesis, we show that their roles are through regulating mitochondria (working model in Fig. 8E). We show that mitochondrial functional and - associated genes’ alterations in *Pkd1*- and *Pkd2*-cKO kidneys begin at the pre-cystic stage. Correcting the defects in ER Ca^2+^ release by *TricB* transgene expression reverses mitochondrial functional perturbations and ameliorates cystogenesis in both *Pkd1-* and *Pkd2*-cKO mice. At least two mechanisms may explain how *TricB* transgene expression reverses mitochondrial dysfunction in both *Pkd1*- and *Pkd2*-cKO. First, PC1 and PC2 form heteromeric Ca^2+^-permeable channels.^29,30^ These heteromeric PC1/PC2 may be a part of ER Ca^2+^ release channels. Second, the carboxyl-terminal tail (CTT) of PC1 is cleaved and translocated to mitochondria to exert anti-cystogenesis.^5,31^ The mechanism of PC1-CTT cleavage and translocation may require normal ER Ca^2+^ homeostasis.

Our study indicates that disruption in ER-mitochondria crosstalk is an early feature in *Pkd*-cKO kidneys. This is illustrated by the global changes in the mitochondrial structure and morphology (decreases in number and size, and shape changes) and as well as disruptions in the close-range ER and mitochondria contact in MAMs. Mitochondrial Ca^2+^ is an essential co-factor for many TCA enzymes.^32^ The close association between IP_3_R in the ER membrane and VDAC in the outer mitochondrial membrane in MAMs is vital for the diffusional transfer of Ca^2+^. Supporting the important role of MAMs in ER-mitochondria Ca^2+^ transfer, mitochondrial Ca^2+^ contents are reduced in *Pkd*-cKO kidneys. Reduction in ER-mitochondrial contact would reduce mitochondrial matrix [Ca^2+^], resulting in reduced ATP production. Beside the inter-organelle transfer of Ca^2+^, ER plays other vital role in the mitochondrial health. The ER is the primary site of lipid synthesis. Lipid transfer from ER is essential for the formation of mitochondrial membranes. Among the mitochondrial lipids, cardiolipin is exclusively synthesized in mitochondria from the ER-imported phosphatic acid. Cardiolipin is crucial for mitochondria cristae formation and directly regulates the function of ETC proteins.^33^ While we have not measured mitochondrial phospholipids, the disruption in ER-mitochondria connection would almost certainly decrease mitochondrial lipids compromising mitochondria membrane biogenesis and function. Overall, the disruption of ER-mitochondria connection would self-amplify leading to further mitochondrial defects and ER-mitochondria disconnection.

Mechanistically, defects in polycystin-regulated ER calcium (Ca²⁺) homeostasis represent a key driver of ER-mitochondria disconnection in *Pkd*-cKO kidneys (Fig. 8E). The expression of the mitochondrial biogenesis master regulator *Pgc1α* is Ca^2+^-dependent.^13^ PGC1α functions as a transcriptional coactivator, binding to the upstream regulatory regions of genes involved in mitochondrial biogenesis and dynamics, including mitofusins (*Mfns*).^34^ Downregulation of *Pgc1α* and its downstream targets, such as *Mfn1* and *Mfn2*, in *Pkd*-cKO mice leads to reduced number of mitochondria, mitochondrial fragmentation and reduced organelle size. Regarding MAMs, MFN2 plays a critical role in the formation of MAMs. Its early downregulation in both *Pkd1*- and *Pkd2*-cKO kidneys, beginning at the pre-cystic stage, likely contributes to reduced MAMs formation. Importantly, these molecular and structural defects are ameliorated by *TricB* transgene expression, supporting the notion that ER Ca²⁺ homeostasis disruption is a central mechanism underlying ER-mitochondria disconnection in *Pkd*-mutant mice

Sigma-1 receptor (S1R) is a Ca^2+^-sensitive ER chaperone regulating ER Ca^2+^ release and ER-mitochondria Ca^2+^ transfer at MAMs.^21^ Besides regulating IP_3_R-mediated Ca^2+^ release, S1R has broader functions on MAMs including promoting lipid exchange, ER membrane remodeling, ER-mitochondrial coupling, and stabilizing MAMs formation.^35^ Defective S1R response contributes to the pathogenesis of neurodegenerative, cardiovascular, and kidney diseases.^35,36^ Pre-084 is a pharmacological S1R activator that promotes MAMs function, and ameliorates diseases in preclinical models.^23^ We found upregulation of S1R expression in *Pkd1*-cKO kidneys, supporting an adaptive response to ER-mitochondria stress. The mere upregulation of S1R is not sufficient to avert cyst formation. Activation of S1R by pre-084 suppresses cystogenesis, supporting the important role of impaired ER-mitochondria communication in the pathogenesis of ADPKD.

In human ADPKD patients, cyst formation starts in the distal earlier than the proximal nephron.^37^ Whether this is due to intrinsic genetic heterogeneities of nephron segments to *Pkd* inactivation or from higher occurrence of somatic second hits in the distal nephron remain unclear. The *Pax8* promoter we used in this study while express higher in distal nephron does express in the proximal as well.^38,39^ In the TEM image analyses, we observed mitochondrial structural changes in proximal as well as the distal tubules. Different from studies of mitochondrial function in *Pkd*-deleted cell lines^12,18,40^, we measured oxygen consumption rate and other functional parameters in manually isolated proximal tubules The requirement of a large number of tubules for accurate measurement limits our experiments to proximal tubules only. The potential quantitative differences in distal tubules cannot be excluded. It should also be mentioned that studies using isolated tubules do not fully recapitulate the *in vivo* setting, perhaps explaining why the differences in the OCR between isolated pre-cystic and control tubules were not as apparent as expected from occurring in the native kidney.

We compared multiple mitochondrial structural parameters between *Pkd*-cKO and control kidneys. The total aggregated differences point to major impacts on the ER-mitochondrial crosstalk. We observed an ∼25% increase in the MAMs distance between *Pkd*-cKO and control (Fig. 2, C and D). The diffusion time (t) is proportional to the square of the distance (γ) (i.e., t ∝ γ^2^). An 25% increase in the distance would lead to 56% ([1.25]^2^ ≈ 1.56) increase in time for Ca^2+^ diffusion across MAMs. Moreover, the length of MAMs covered by the ER membrane is also reduced. At the global level, the reduction in size and number of mitochondria would further add to the disruption in the ER-mitochondria crosstalk. The independent measurement by PLA that assesses a broader picture of the ER-mitochondria connectivity shows ∼80% reduction in the interaction. Together, our results provide compelling evidence that a functionally significant disconnection between the ER and mitochondria occurs prior to cyst formation. The reason why cysts take weeks to form following the disruption of the ER-mitochondria crosstalk in *Pkd*-mutant mice remains unknown. Additional processes, as well as the loss of polycystin function in other subcellular compartments, are likely involved.

Epigenetic changes occur in ADPKD.^26^ How *PKD* mutations lead to epigenetic rewiring is not fully understood. Increased glycolysis with elevated cellular acetyl-CoA levels accompanies mitochondrial dysfunction in ADPKD.^5,18,26^ An increase in the nuclear acetyl-CoA pool promotes histone acetylation for gene activation.^41^ Lakhia *et al.*, recently reported histone acetylation at enhancers and super-enhancers is important for the upregulation of *cMyc* in *Pkd1*-mutant mice.^26^ Here, we show that ER-mitochondria disconnection is an underlying mechanism of these epigenetic remodeling. We show that increased histone acetylation at *cMyc* enhancers occurs in the pre-cystic and cystic kidneys of both *Pkd1*- and *Pkd2*-cKO mice and reversed by *TricB* expression or treatment with pre-084. CDK7 is a transcriptional kinase and essential component of the transcription factor-II H (TFIIH) involved in the transcription initiation and DNA repair.^42^ *Cdk7* is upregulated in ADPKD and plays a pivotal role in organizing and stabilizing the super-enhancer complex for genes involved in metabolic reprogramming.^10^ We further show that increased histone acetylation of *Cdk7* enhancers occurs in *Pkd1*- and *Pkd2*-cKO beginning at the pre-cystic stage and reversed by *TricB* expression or treatment with MAMs activator pre-084. Thus, metabolic reprogramming in ADPKD occurs, at least in part, through the loss of *PKD* function in the ER leading to mitochondrial stress and mitochondria-driven epigenetic rewiring.

In summary, defects in polycystin function in the ER contribute to mitochondrial perturbations in ADPKD. Mitochondrial dysfunction fuels epigenetic shifts contributing to widespread gene rewiring. Notable downstream events include activation of *cMyc* to promote cell proliferation and *Cdk7* to promote metabolic reprogramming. These events, alongside defective polycystin function at the cilium, plasma membrane, and other subcellular compartments, converge to initiate and sustain cystogenesis. Restoring ER-mitochondria communication may represent a promising therapeutic strategy for ADPKD.

## Materials and Methods

### Animal Usage

*Pkd2-cKO* (*Pkd2-flox/flox;Pax8-rtTA;TetO-Cre*) and *Pkd1-cKO* (*Pkd1-flox/flox;Pax8-rtTA;TetO-Cre*) mice were obtained from the Maryland PKD Core Center (https://www.pkd-rrc.org/pkd1-pkd2-pkhd1-rodent-models/). All animal maintenance and experiments were conducted in accordance with the Guide for the Use and Care of Laboratory Animals and approved by the Institutional Animal Care and Use Committee. Pkd2 and Pkd1 conditional knockout in kidney were induced by 2 mg/ml doxycycline in 2% sucrose in drinking water or by a 0.625g/kg doxycycline diet (TD.240234) for 2 weeks, starting at the age of 4 weeks. Controls mice were littermates lacking either Pax8-rtTA or Tet-O-Cre, or both. They were given doxycycline. Mice are of C57BL/6 genetic background. Transgenic mouse *TricB* (*cTg_Tric-b*) was generated as described before.^6^ Genotyping primers and protocol are as previously described (https://www.pkd-rrc.org/pkd1-pkd2-pkhd1-rodent-models/).^6^ Except where indicated otherwise, male mice were used to avoid the possibility of sexual dimorphism.

### Histology, cyst Index, and blood chemistry analysis

Kidneys were fixed with 10% formalin or 4% PFA, paraffin embedded, sectioned, and stained by hematoxylin and eosin. The cyst index was analyzed with CystAnalyzer.^43^ For measurement of BUN, serum was collected from euthanized animals and was analyzed using a BUN assay kit (Pointe Scientific B7552150).

### Western blotting

Total kidney lysates were prepared by homogenization in RIPA buffer (Thermo Scientific) supplemented with protease inhibitor cocktail (Roche). Total protein concentration was measured by Bradfords (Bio-Rad, 5000006) or Pierce™ BCA Protein Assay Kits (Thermo Fisher Scientific 23225). Protein lysates were boiled in 1× NuPAGE LDS Sample Buffer with 10mM DTT, loaded on to NuPAGE 4% to 12% Bis-Tris Gels (Thermo Fisher Scientific) and run at 130 V. Proteins are then transferred to 0.45 µm PVDV membrane (Thermo Fisher Scientific) at constant 300 mA for 80 minutes at 4°C. The membrane was blocked in 5% skim milk in TBS-Tween 20 (0.01%) for 60 min at RT and incubated in following primary antibodies overnight at 4°C; PC2 (sc-28331, 1:500; Santa Cruz Biotechnology), VDAC (600-101-HB2, 1:1000, ThermoFisher Scientific), S1R (PA5-30372, 1:500, ThermoFisher Scientific), IP3R (PA1-901, 1:1000, ThermoFisher Scientific), Calnexin (sc-6465, 1:1000, ThermoFisher Scientific), TricB (HPA018465, 1:500, Sigma). Membranes are then washed three times in 1x TBST each for 10 min and incubated in following HRP-conjugated secondary antibodies for 1hr at RT; anti-Mouse HRP (7076P2, 1:10000, Cell Signaling), anti-Rabbit HRP (G21234, 1:10000, ThermoFisher Scientific), anti-Goat HRP (sc-2352, 1:10000, Santa Cruz Biotechnology). Afterwards, membranes are washed three times in TBST for 10 min each and developed in Clarity Western ECL Substrate (BioRad, 1705061) or SuperSignal™ West Femto (ThermoFisher Scientific, 34096X4). Signal is then detected in ChemiDoc XRS+ System with Image Lab Software (Bio-Rad).

### Real-time PCR

Kidney tissues were dissected and saved in RNAlater. Total RNA from kidney tissues or microdissected proximal tubules were isolated by homogenization in TRIzol (ThermoFisher Scientific, 15596026) following manufacturer’s instructions. Subsequently, cDNA was synthesized from 1μg of RNA by using iScript cDNA Synthesis Kit (Bio-Rad, 1708891). Primers were designed through Primer-BLAST and procured from IDT (Supplementary Table 1). 18s rRNA or GAPDH gene expression was used as an endogenous control for normalization of targeted gene expression. Quantitative real-time PCR was then performed using iTaq Universal SYBR Green Supermix (Bio-Rad, 1725124) in the C1000 Touch Thermal Cycler (Bio-Rad) following the manufacturer’s instructions. ΔΔCt method was used to express the fold difference of targeted gene expression.

### Mitochondrial membrane potential assay

Mitochondrial membrane potential is measured by using cell permeant fluorescent dye, Tetramethylrhodamine, methyl ester (TMRM) (ThermoFisher, T668) following manufacturers protocol and McKezie *et al*.^44^ Briefly, PTs were isolated as described elsewhere from dissected kidneys and seeded on glass cover slips coated with Cell-Tak (Corning™) and cultured in complete media (Lifeline, RenaLife ™ Epithelial Medium Complete LL-0025) for 72 hr at 37°C in the CO_2_ incubator. On the day of experiment, seeded cells were incubated at 37°C for 30 min with TMRM (100 nM) in the cell growth media along with MitoTracker Green (200 nM) (ThermoFisher, M7514), a mitochondrial-selective dye that is used to track mitochondria.

### Mitochondria and MAMs isolation

Mitochondria and MAMs were isolated from kidney following Wieckowski *et al*.^45^ Briefly, both kidneys were homogenized in the homogenization buffer (225 mM mannitol, 75 mM sucrose, 0.5% BSA, 0.5 mM EGTA and 30 mM Tris–HCl at pH 7.4) using Dounce homogenizer and then centrifuged at 740g for 5 min to remove unbroken cell debris. Collected supernatant again centrifuged at 9,000g for 10 min at 4°C and the pellet containing mitochondria sequentially washed as per recommended protocol. Finally, crude mitochondrial pellet is resuspended in MRB (mitochondria resuspending buffer: 250 mM mannitol, 5 mM HEPES and 0.5-mM EGTA at pH 7.4) and layered on top of percoll medium (225 mM mannitol, 25 mM HEPES, 1 mM EGTA and 30% Percoll at pH 7.4) followed by MRB solution and centrifuged at 95,000g for 30 min at 4°C in an ultracentrifuge (Beckman Coulter, Optima XPN-100 Ultracentrifuge). Following centrifugation, floating diffused white band representing MAM and dense band at the bottom for mitochondria were collected and rewashed accordingly to obtain pure mitochondria and MAM in MRB solution. The purity of mitochondria and MAM preparation was checked by western blot analysis of targeted markers.

### Mitochondrial calcium measurement

Measurement of Ca^2+^ concentration in the mitochondria of cultured primary proximal tubule cells were performed following Maxwell *et al*.^20^ Accordingly, Rhod-2 AM (20 μm) (ThermoFisher, R1244) is used as a fluorescent labeled calcium indicator and MitoTracker green (200 nM) in Tyrode’s solution (140 mM NaCl, 5 mM KCl, 1.2 mM CaCl2, 1 mM MgCl2, 0.33 mM NaH2PO4, 5.5 mM glucose, 10 mM HEPES at pH 7.4) for 30 min at room temperature in dark. Afterwards, de-esterification of Rhod-2 AM is done by replacing the dye solution with fresh Tyrode’s solution and incubated for another 30 min. Cover slips were then transferred to imaging chambers and plasma membrane of cells were permeabilized by 0.005% saponin solution for 1 min to offload cytosol-localized Rhod-2 AM while retaining mitochondria localized Rhod-2 AM. Buffer in the microscope chamber is then immediately replaced by Ca^2+^-free Tyrode’s solution with 2 mM EGTA. Subsequently, fluorescent images showing colocalization of Rhod-2 AM (Ex/Em 552/581 nm) and MitoTracker Green (Ex/Em 490/516 nm) were taken under the microscope (Nikon Eclipse E600) and processed in ImageJ (1.54f).

Absolute Ca^2+^ content was measured in isolated mitochondrial fractions (see mitochondria isolation method) using colorimetric Calcium assay kit from Abcam (ab102505) and following Kwong *et al*.^46^ Briefly, mitochondrial pellets were resuspended in calcium assay buffer supplied in the kit and solubilized using a sonicator. Afterwards, the lysates were centrifuged at 10,000 rpm for 3 min at 4°C to clear insoluble material. Protein quantification was done through BCA assay, and lysates were loaded onto wells followed by chromogenic reagent and calcium assay buffer provided in the kit. After mixing and incubating in the dark at room temperature for 5 min, the absorbance is measured at 570 nm using a microplate reader. Observed OD is then used to calculate the absolute Ca^2+^ content by using a series of Calcium standards supplied in the kit.

### Measurement of total H3K27ac

Total H3K27ac was measured in mice kidney tissues by using EpiQuik Global Acetyl Histone H3K27 Quantification kit (Epigentek, P-4059) and following Lakhia *et al*.^26^ Histone extracts were prepared through acid-extraction method following manufacturers protocol. Kidney tissues were minced and disaggregated in TEB (PBS containing 0.5% Triton X 100, 2 mM PMSF and 0.02% NaN_3_) buffer using a Dounce homogenizer. Tissue homogenate is then centrifuged at 10,000 rpm for 1 min at 4°C and the collected tissue pellet after resuspending in 3 volumes of extraction buffer (0.5N HCl + 10% glycerol) incubated on ice for 30 min. After centrifugation at 12,000 rpm for 5 min at 4°C, the supernatant is collected, and 8 volumes of acetone is added and incubated for overnight at -20°C. Afterwards, it is centrifuged again at 12,000 rpm for 5 min, air-dried the pellet and dissolved in nuclease free water. Protein extracts were then quantitated through BCA protein assay. Following manufacturers protocol, 200 ng of histone extracts were added per well for H3K27ac quantification. Briefly, acetylated histone H3 at lysine 27 from the extracts were captured onto the strip wells coated with anti-acetyl H3K27 antibody. Captured H3K27ac were then detected through a colorimetric assay with the help of a labeled detection antibody specific to H3K27ac and the absorbance is measured in a microplate reader at 450 nm. Using a reference standard control supplied in the kit the absolute amount of H3K27ac is calculated.

### ChIP-qPCR

ChIP-qPCR assays were carried out using rabbit Anti-Histone H3 (acetyl K27) antibody (Abcam ab4729) following manufacturer’s protocol (Santa Cruz). Briefly, tissues were cut into small pieces (∼1mm^3^) and cross-linked by 1.5% formaldehyde at RT for 15 min. Following quenching by 0.125M glycine tissues were washed twice in ice-cold PBS. After tissue homogenization by Dounce tissue homogenizer, cells were lysed in lysis buffer (Santa Cruz sc-45000) and nuclei pellets were collected by centrifugation and washed once with ice-cold PBS. Pellets were resuspended in high salt lysis buffer (Santa Cruz, sc-45001) and sonicated on ice 30 times at 30 sec intervals. After centrifugation of the sonicated sample supernatant chromatin was collected and quantified. For immunoprecipitation, Protein A/G Plus-Agarose beads (Santa Cruz, sc-2003) were used to preclear as well as for IP. Precleared chromatin samples were incubated with H3K27ac antibody overnight at 4^0^C followed by incubation with agarose beads for 2 hrs at 4^0^C. Beads were then collected and washed twice with high salt lysis buffer followed by wash buffer four times (Santa Cruz, sc-45002). Following elution in elution buffer (Santa Cruz, sc-45003) chromatin samples were reverse-crosslinked in a water bath at 67^0^C and then extracted by phenol-chloroform purification method. Targeted genomic loci with H3K27ac marks were then quantified by designed primers through quantitative real-time PCR as above (Supplementary Table 2) and represented as percentage input following the expression Input % = 100/2^ΔCt^ ^[normalized^ ^ChIP]^. Thirteen loci with H3K27ac marks surrounding *Cdk7* gene were selected as shown by different cell type class in ChIP-Atlas, visualized in integrative genomics browser (IGV2.18.4) and UCSC genome browser. For *cMyc*, 3 sites (E5, E19 and E28 corresponding to a previously reported study were chosen.^26^

### Cell culture and CRISPR/Cas9 genome editing

Mouse distal convoluted tubule (MDCT) cells were cultured in Dulbecco’s Modified Eagle Medium, 10% FBS, 100 U/ml Pen Strep and maintained at 37°C and 5% CO2. These cells were used to examine the enhancer effect of C5 and C10 loci on *Cdk7* gene expression. For deletion of genomic regions surrounding C5 and C10 a pair of sgRNA (SgRNA1 and 2, Supplementary Table 3) was designed (CRISPR-Cas9 gRNA checker) for each site and cloned into PX459 (Addgene plasmid 62988) construct. MDCT cells were then transfected by CRISPR constructs and 36 hr post transfection selected by puromycin (2 μg/ml) for additional 48 hr. Cells were then analyzed for homozygous deletion by genotyping primers (Supplementary Table 3). Genomic region of 1,301 bp and 1,889 bp surrounding C5 and C10 loci were deleted (Figure 8D). Homozygous deleted cells were then used to check the *Cdk7* gene expression and compared to control cells.

### Chromatin conformation capture assay (3C)

3C assays were performed following Hagege *et al*. and Tolhuis *et al*.^47^ Fixed chromatin samples were prepared from kidney tissue lysates following tissue mincing (as described above) and subjected to formaldehyde crosslinking and cell lysis. Restriction enzyme BamH1 (NEB, R3136S) that generates 5’ overhangs was selected as the enzyme of choice based on the proximity to the *Cdk7* promoter as well as to putative enhancers with H3K27ac marks. Cross-linked chromatins were digested overnight at 37^0^C with continuous shaking. Following digestion and dilution with T4 DNA ligation buffer chromatin samples were ligated by T4 DNA ligase (NEB) overnight at 16^0^C. Following de-crosslinking by proteinase K at 65^0^C, the ligated chromatin samples were purified by phenol-chloroform extraction method and concentrated. Sample purity of the purified chromatin samples were checked as per Hagege *et al*.^47^ Loading adjustments were done by using an internal primer set in mice *Gapdh* gene containing no BamH1 restriction site. In total, 21 BamH1 restriction sites were chosen; 7 each centered around the targeted loci of C1, C5 and C10 and 1 BamH1 restriction site close to the *Cdk7* promoter region (Supplementary Table 4). Additionally, for generation of control template to normalize primer efficiency across different primer sets, genomic regions surrounding each restriction site are individually amplified by PCR, BamH1 digested, purified and ligated in equimolar amount. The serial dilution of this PCR control template was used to generate standard curves to calculate the slope (a) and the y-intercept (b) for each primer set. Afterwards, prepared 3C chromatin samples were used to calculate the ligation or relative crosslinking frequency by qPCR using different 3C primers tagging each restriction fragment against the constant or anchor fragment on the promoter of Cdk7 gene following the expression =10^(Ct–b)/a^. Further, the results were normalized to the control interaction frequencies between two restriction fragments located in *Ecc3* gene as per Hagege *et al*.^47^

### Proximity ligation assays

Proximity ligation assays were performed using the kit Duolink® In Situ Red Starter Kit Goat/Rabbit (Sigma DUO92105). Steps were followed according to the manufacturer’s protocol. Briefly, mice kidney tissues were fixed in 4% formaldehyde, embedded in OCT and 6 µm sections were cut through a cryostat. Tissue sections on slide were rinsed twice in water, washed in TBS plus 0.025% Triton X-100 twice for 5 min each and then blocked using Duolink® Blocking Solution for 60 minutes at 37 °C in a humidity chamber. After incubation, blocking solution is drained and Duolink® Antibody Diluent containing primary antibodies specific to IP3R1 (Thermofisher PA1-901, 1:250 dilution) and VDAC (Thermofisher, 600-101-HB2, 1:250 dilution) were added and incubated overnight at 4°C. Slides were then washed in 1x wash buffer A twice for 5 min each at RT and incubated in PLA probe solution in a humidity chamber for 1 hour at 37^0^C. Subsequently, slides were washed again with 1x Wash Buffer A twice for 5min each at RT and incubated with ligation solution for 30 minutes at 37 °C. Following washing with 1x wash buffer A twice, slides were incubated with amplification solution in a pre-heated humidity chamber for 100 minutes at 37 °C. Afterwards, slides were washed twice with 1x wash buffer B for 10 min each followed by 0.01x wash buffer B for 1 min. Sections were then mounted with Duolink® In Situ Mounting Medium with DAPI and visualized on Leica SP8 confocal microscope (Leica) through LAS X 3.0.14 software platform (Leica) with a 100x objective. At least three images from each mice were processed and mean fluorescence intensity were analyzed in ImageJ 1.54f.

### Microdissection of proximal renal tubules

Proximal tubules were dissected and isolated following Glaudemans et al.^17^ Briefly, mice kidneys were dissected, sliced, and incubated in prewarmed Hank’s balanced salt solution (137mM NaCl, 5mM KCl, 0.8mM MgSO4, 0.33mM Na2HPO4, 0.44mM KH2PO4, 1mM MgCl2, 10mM Trishydroxtmethyl aminomethane hydrochloride, 0.25mM CaCl2, 2mM L-glutamine, 2mM L-lactate, 295 osmolality at pH 7.4) containing collagenase type I (1.5 mg/ml) and bovine serum albumin (1mg/ml) and shaken vigorously on a titer plate shaker at 37^0^C for 30 min. Following incubation, the digested segments were passed through 200 µM (pluriStrainer 200µm) filter and collagenase digested tissue segments were collected. Microdissection of individual proximal renal tubules were then performed under a stereo microscope based on their morphological characteristics following Glaudemans et al.^17^

### Oxygen consumption rate assay

The measurement of oxygen consumption rate (OCR) is carried out according to manufacturer’s protocol (Agilent Seahorse XF Cell Mito Stress Test Kit). Individually isolated 8-10 proximal renal tubules were seeded on a coated (Corning™ Cell-Tak) Seahorse XFp Cell Culture Miniplates (Seahorse XFp FluxPak, Agilent) with Seahorse XF DMEM media supplemented with 1 mM pyruvate, 2 mM glutamine, and 10 mM glucose and incubated in a non-CO_2_ incubator for 1hr at 37^0^C. For PRE-084 treatment, media is also supplemented with 1 µM of PRE-084 (Bio-techne, 0589). Following incubation, media is replaced with fresh media and OCR is measured in the seeded tubules using Seahorse XF Cell Mito Stress Test Kit (Agilent) in a Seahorse XF HS Mini (Agilent) as per manufacturer’s instructions. Preoptimized concentrations of modulators provided in the kit were used, oligomycin (1.5 µM), FCCP (1 µM) and rotenone and antimycin A (0.5 µM). Each OCR assay is performed in 2-3 technical replicates per group. After OCR measurement, total RNA from the seeded tubules was extracted by Trizol and gene expression of 18s rRNA were quantitated using qRT-PCR and cycle threshold (Ct) values were used for normalization across wells. Data were analyzed by Seahorse XF Mito Stress Test Report Generator.

### Transmission and scanning electron microscopy

Structural analysis of mitochondria and MAMs in kidney tissue samples were analyzed as per Lam *et al*. 2021 and Giacomello *et al*. 2016 using ImageJ v1.53t.^9,48^ For visualization, TEM images were taken at 10k magnifications. For mitochondrial morphology analysis, random TEM images were taken from multiple embedded sections derived from each mice sample at 10k magnification. Subsequently, each image was divided into quadrants by using ImageJ plugin quadrant. After splitting into four quadrants, two quadrants were randomly selected for mito and MAMs structural analysis. To ensure an unbiased approach, selected quadrants from all samples were anonymized and pooled, and the rater conducting the analysis was blinded to the experimental groups. Randomly, 30 mitochondria from multiple images were included for analysis per sample. MAMs characterization was performed based on previous studies.^9,48^ Within the mitochondria, only those MAMs were analyzed where the ER-mito contacts distance was less than 30 nm.^9^ At least 25 mitochondria were selected randomly from multiple images per group for measurement of MAMs distance and MAMs length in relation to mitochondrial perimeter.

### Statistical analysis

Quantitative data are presented as mean ± SEM. Statistical significance was analyzed by one-way or two-way ANOVA followed by Tukey’s or Sidak’s multiple comparison tests or by two-tailed, unpaired Student’s t test in GraphPad Prism (Version 10.2.0), as appropriate. All experiments were repeated at least three times. p < 0.05 was considered as statistically significant.

## Funding and Acknowledgements

We thank University of Iowa Central Microscopy Research Facility for TEM studies and Dr. Jianqiang Shao for advice. This work was supported in part by NIH (DK-109887) and philanthropic gifts from Jared and Carol Hills.

