## Supplemental file with 6 supplementary figures and 4 supplementary tables for "Endoplasmic reticulum-mitochondrion disconnection promotes metabolic reprogramming and cystogenesis in polycystic kidney disease" (from another version of this preprint: version 1, 2025-11-01)

The file contains 6 supplementary figures & 4 supplementary tables

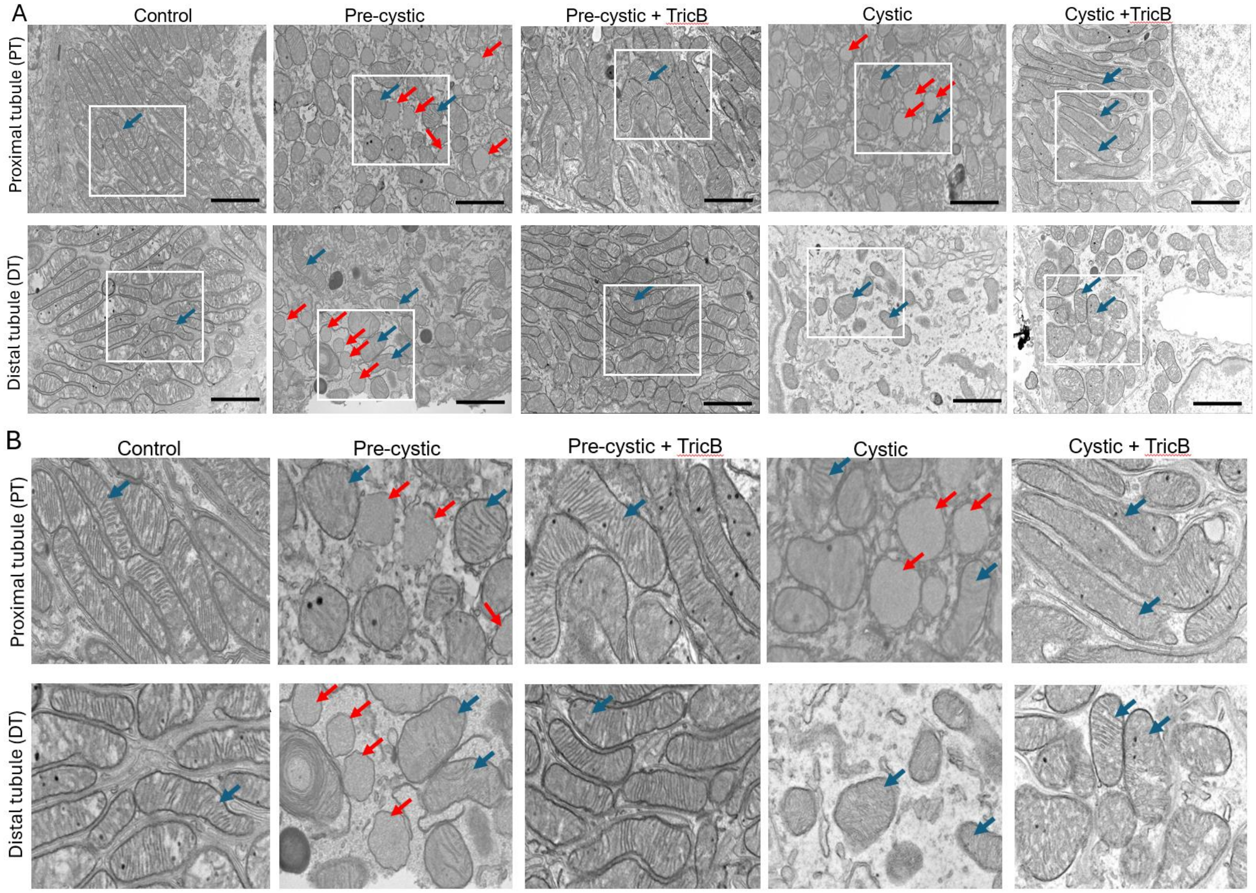


**Supplementary Figure 1. TEM images of mitochondria in *Pkd2*-cKO mice.** (**A**) Representative TEM images at 10K magnification illustrating mitochondrial morphology and cristae structure in control, *Pkd2*-cKO mice at pre-cystic and cystic stages with or without *TricB* expression. Upper row- Proximal tubules (PT) and Lower row- Distal tubules (DT). Scale bar 2µm. (**B**) Enlarged images from white boxed area above showing distinct cristae in control and *TricB* expressing mice in *Pkd2*-cKO mice. Compared to the controls, mitochondria in *Pkd2*-cKO are fewer and more rounded in shape, and with decreased cristae density but with increased numbers of lysosomes. Upper row- Proximal tubules (PT) and Lower row- Distal tubules (DT). Blue and red arrow indicates mitochondria and lysosomes, respectively. Mitochondria are identified by cristae and dense membrane (2 layers of lipid bilayer- outer and inner mitochondria membrane). Lysosomal membrane contains only 1 layer of lipid bilayer and no cristae.

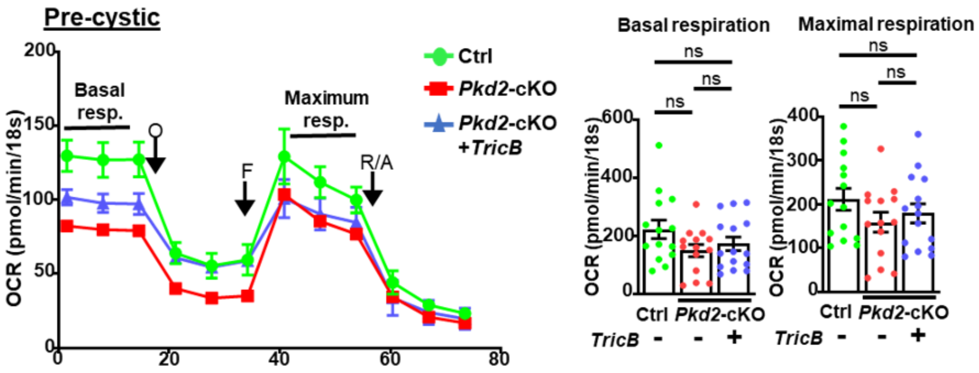


**Supplementary Figure 2.** OCR from PTs of control and pre-cystic *Pkd2*-cKO kidneys with or without *TricB* expression. Assessed by one-way ANOVA with Tukey's multiple comparison test. ns- not significant.

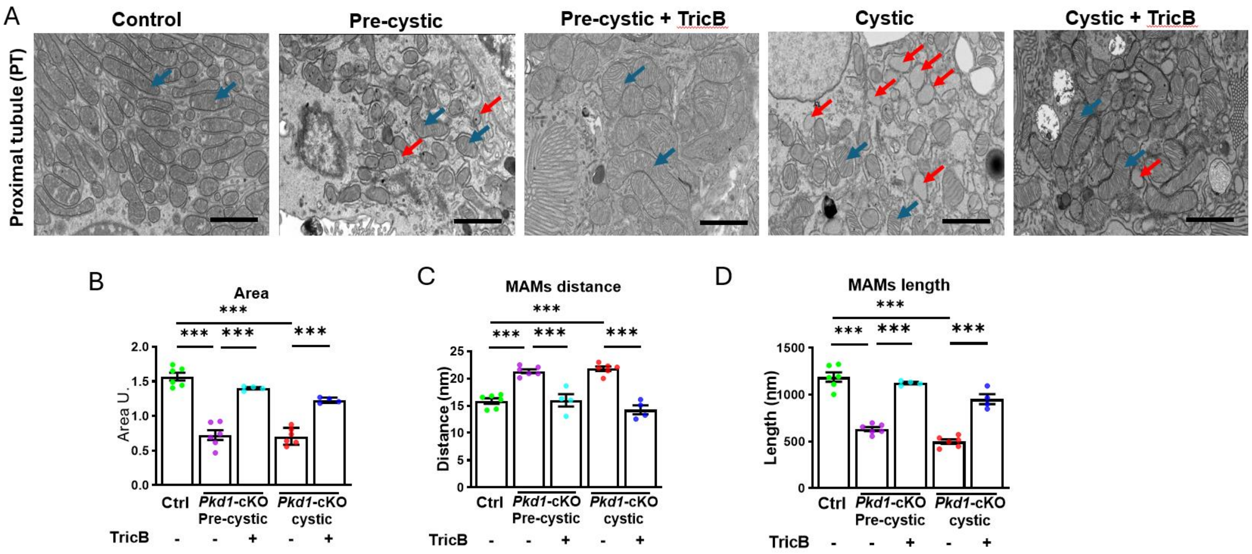


**Supplementary Figure 3. Structural analysis of mitochondria and MAMs in *Pkd1*-cKO and reversal by TricB.** (**A**) Representative TEM images at 10K magnification illustrating mitochondrial morphology and cristae structure in control and *Pkd1*-cKO mice at pre-cystic and cystic stage with or without TricB expression. Blue and red arrow indicates mitochondria and lysosomes, respectively. (**B**-**D**) Average mitochondria area (**B**), MAMs distance (**C**), and length (**D**) in *Pkd1*-cKO mice at the pre-cystic and cystic stage with or without *TricB* expression in comparison to control. Each data point is average of total 30 mitochondria from multiple images per mouse. 6 mice per group were analyzed. Data are mean ± SEM. Assessed by one-way ANOVA with Tukey's multiple comparison test. *, **, ***; $p < 0.05$, 0.005, 0.001, respectively. ns- not significant.

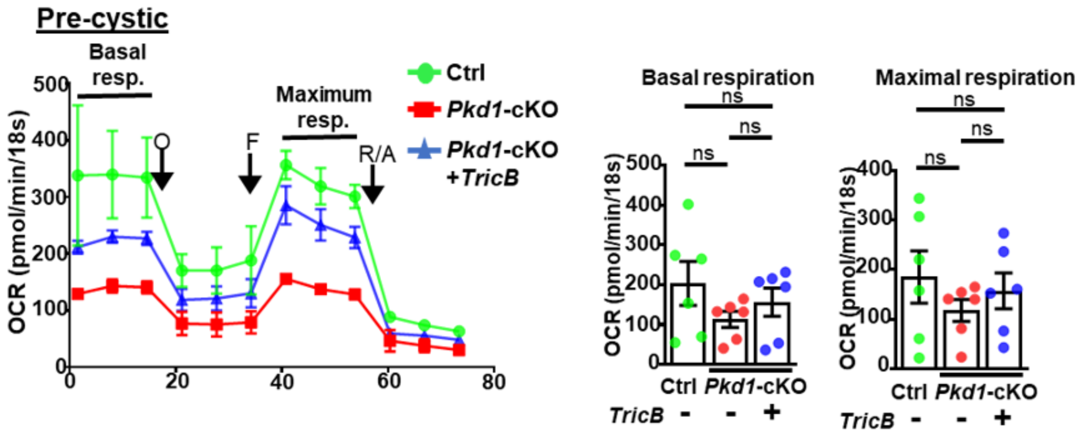


**Supplementary Figure 4.** Oxygen consumption rate (OCR) in PTs freshly isolated from control, *Pkd1*-cKO with or without *TricB* expression at pre-cystic stage. Left panel shows representative line tracings of one experiment. Data points in each group represent mean ± SD of multiple wells from a single mice per group with each well containing 9-10 freshly dissected PTs. OCR values were normalized to 18S rRNA measured by rt-PCR. Bar graphs on the right represent mean ± SEM of 6 experiments. Basal respiration is OCR before addition of ATP synthetase inhibitor oligomycin. Maximal respiration is OCR after addition of mitochondrial OXPHOS uncoupler FCCP. Assessed by one-way ANOVA with Tukey's multiple comparison test. ns- not significant.

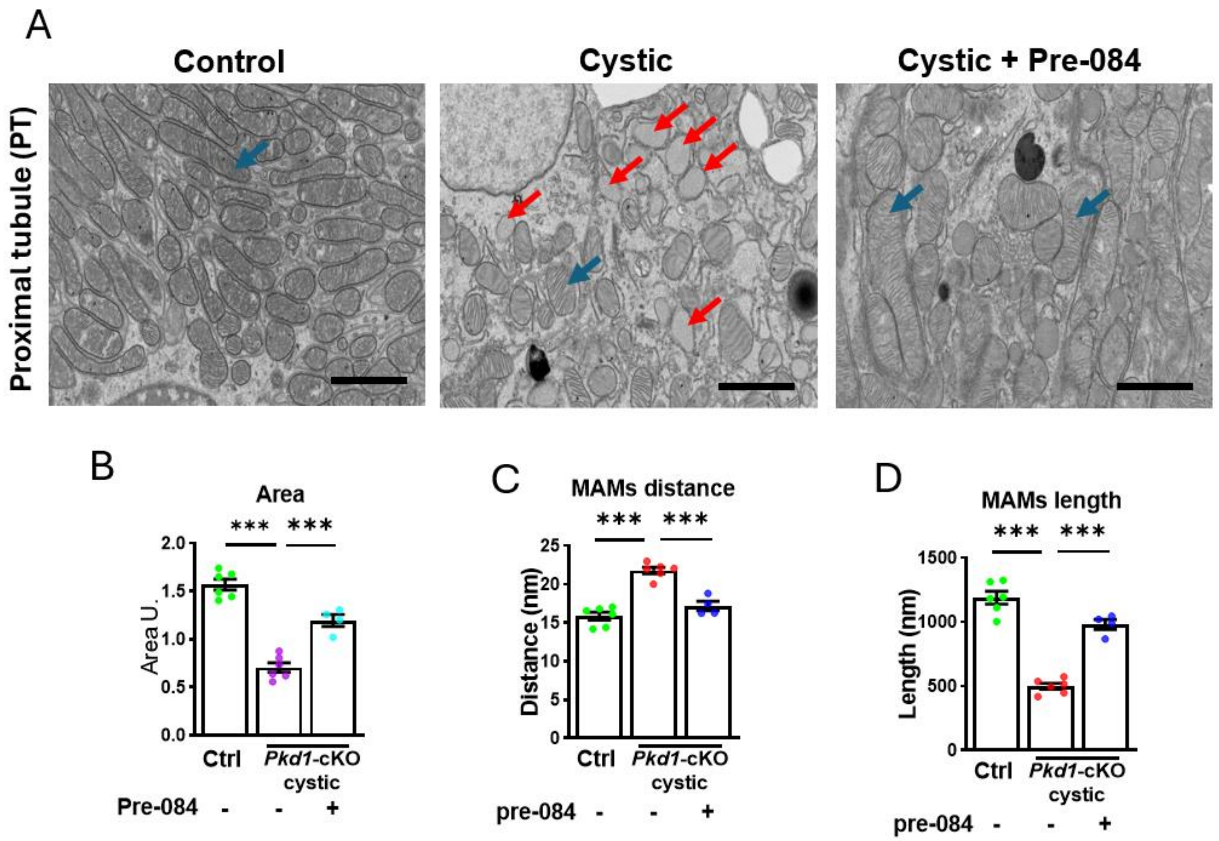


**Supplementary Figure 5. Structural analysis of mitochondria and MAMs in *Pkd1*-cKO and reversal by pre-084.** (**A**) Representative TEM images at 10K magnification illustrating mitochondrial morphology and cristae structure in control and *Pkd1*-cKO mice at cystic stage with or without pre-084 treatment. (**B**-**D**) Average mitochondria area (**B**), MAMs distance (**C**) and length (**D**) in *Pkd1*-cKO mice at the cystic stage with or without pre-084 treatment in comparison to control. Each data point is average of total 30 mitochondria from multiple images per mouse. 6 mice per group were analyzed. Data are mean ± SEM. Assessed by one-way ANOVA with Tukey's multiple comparison test. ***, $p < 0.001$.

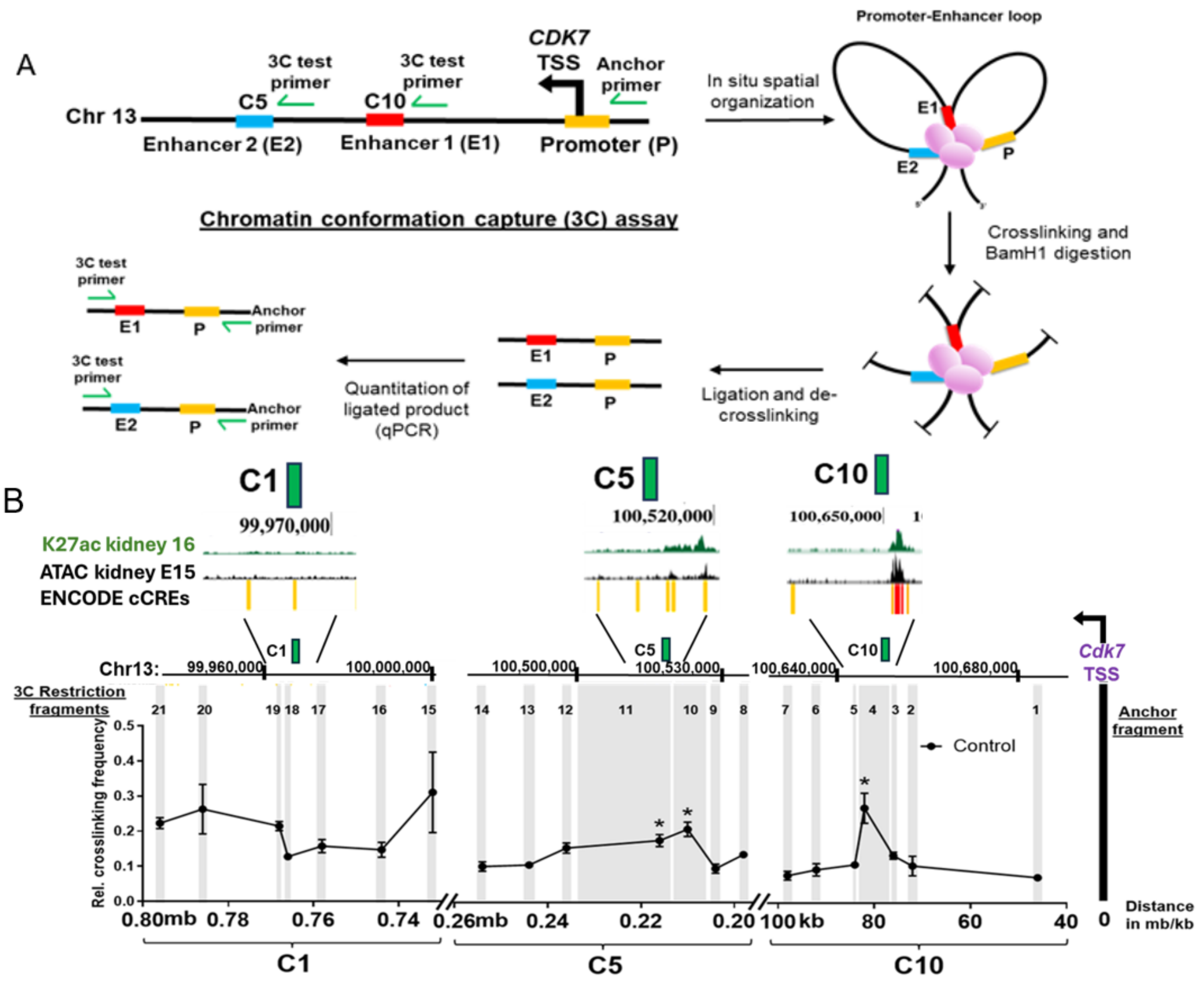


**Supplementary Figure 6. H3K27ac-enriched C5 and C10 genomic regions are *Cdk7* enhancers.** (**A**) Diagrammatic representation of chromosome conformation capture (3C) assays designed to validate chromatin-chromatin interactions between potential H3K27ac-enriched genomic regions near the *Cdk7* gene and the *Cdk7* promoter. Each 3C primer targets nearby chromatin regions, referred to as 3C restriction fragments, while the constant anchor primer near the *Cdk7* TSS (transcription start site) serves as the anchor fragment. For quantification of each prospective ligated product, the anchor primer was used in combination with each respective 3C primer. (**B**) Top: enlarged illustration of H3K27ac marks depicted in the UCSC genome browser aligned with database of "K27ac kidney 16", "ATAC kidney E15", and "ENCODE cCREs". K27ac kidney 16: kidney acetylated histone K27; ATAC kidney E15: "assay for transposase accessible chromatin" kidney E15; ENCODE cCREs: ENCODE putative cis-regulatory elements. Relative crosslinking frequencies between targeted restriction fragments (gray vertical bars) surrounding C1 (21-15), C5 (14-8) and C10 (7-1) putative enhancers with that of anchor fragment (black vertical bar to the right) located nearby *Cdk7* TSS in prepared 3C chromatin samples from control mice kidneys. Data are presented as mean ± SEM from 3 mice per group. Unpaired student t-test; *; $p < 0.05$ for fragment 10,11 vs fragment 8, 9, 13, 14 for C5; fragment 4,5 vs fragment 1, 2, 3, 6, 7 for C10, by 2-way ANOVA with post-hoc Tukey test.

**Supplementary Table 1. Primers used for qRT-PCR analysis of mitochondria regulatory genes**

| Name | Sequence |
|---|---|
| ND1 | CTA GCA GAA ACA AAC CGG GC |
| | CCG GCT GCG TAT TCT ACG TT |
| HK2 | GCC AGC CTC TCC TGA TTT TAG TGT |
| | GGG AAC ACA AAA GAC CTC TTC TGG |
| PPARa | TGG TGC ATT TGG GCG TAT CT |
| | CAC AGA GCG CTA AGC TGT GA |
| PGC1a | GTG CTG TGT GTC AGA GTG GA |
| | CAA CCA GAG CAG CAC ACT CTA |
| Mfn1 | TTTGGCTGATCGGTGTACCA |
| | CGAACACAGCTTGTGGCAAT |
| Mfn2 | CAA GAC CGG CTG AGG TTT ATT |
| | CCT TTC CAC TTC CTC CGT AAT C |
| Drp1 | CCAGAGGAACTGGTGTGGTC |
| | TTTTCCATGTGGCAGGGTCA |
| Opa1 | CACTGGCAAACGAGTTCAGC |
| | TGAGCAGGATTTTGACACAGTCT |
| Cdk7 | GGTCTGGTTTCTCACTCGCA |
| | TAGACCGTCGCAAACTGTCC |

**Supplementary Table 2. Primers used for ChIP-qPCR analysis of H3K27ac-enriched genomic marks**

| Name | Sequence |
|---|---|
| C1F | CAGCTTGGTTTGTTCACGGG |
| C1R | AAATGGTGTGACTGCCCTCG |
| C2F | TTTTGAAGCCAGGGGTGTGT |
| C2R | ATCCAAGGGCAACACTTCGT |
| C3F | ATCGAGCCTGTCCGCTCT |
| C3R | AGTATGGCTGCGGGGTCG |
| C4F | GCTTTCATTTGTGCAAGTTACCA |
| C4R | TATCAACAAACAGGATAGGTGGCT |
| C5F | GGCCTATTGTTTTGTGCTATCCT |
| C5R | ATTAAATCTGAAAGGACAATGGCA |
| C6F | GGTTCAACCCCGACCGAGTA |
| C6R | TGGAATGAGCTTGGGTAGGC |
| C7F | GTTCGGCCTATTGTCTCCACA |
| C7R | TGTTGTGACTTCGGTCTTGGT |
| C8F | TGAGTACTATAGTCAGCCCCTCA |
| C8R | TTTCGATGACAACGACGGGT |
| C9F | AGAGTGCACCCTTAATACCACCTA |
| C9R | CATGCCCAATTTTAGCCATCTAGT |
| C10F | TGGCTAGCTTAACCACAGGC |
| C10R | GGCTCCCAGTTCTCTTGCAT |
| C11F | TCTGAGGCCGTGTCTTTGTC |
| C11R | CTCCCGTGTTCTGCATCCAT |
| C12F | GCACAGCATGAATGCGAGAG |
| C12R | ACAAAAGGATCCCGCCAGAG |
| C13F | GTGGTAGCACTCGCCTATACAT |
| C13R | AAAGGATTAAAGGCATGTGTGATT |

**Supplementary Table 3. Oligos used for CRISPR/Cas9 mediated deletion of C5 and C10 surrounding regions**

| Name | Sequence |
|---|---|
| C5_sg1_sen | CACCGGAGAACTTGTGCAAGGATCG |
| C5_sg1_antisen | AAACCGATCCTTGCACAAGTTCTCC |
| C5_sg2_sen | CACCGGACTGTCTGAACCACCCAGG |
| C5_sg2_antisen | AAACCCTGGGTGGTTCAGACAGTCC |
| C10_sg1_sen | CACCGATTATGGCCTGACCTTTCAA |
| C10_sg1_antisen | AAACTTGAAAGGTCAGGCCATAATC |
| C10_sg2_sen | CACCGTTGGTAGACTACTCACTAGT |
| C10_sg2_antisen | AAACACTAGTGAGTAGTCTACCAAC |
| C5_GenoF | TCCTCACACTCACCACACTC |
| C5_GenoR | GATAGCCCTTCACTCTCCCC |
| C10_GenoF | TGTCATGCTACCAAGTACTGTG |
| C10_GenoR | GGTTGGCCTTTGACTTCTCA |

**Supplementary Table 4. Primers used for qRT-PCR for chromatin conformation capture (3C) assays**

| Name | Sequence |
|---|---|
| 3C_CDK7 | GGCAGCCTCCCCTTTTCATT |
| Control_CDK7F | TGGTTGAGCTTCGTTCCTCT |
| Control_CDK7R | AAGCAAGGTCTCAGTCCCTG |
| 3C_C1_B1 | CCCTGTGGTTACACCCATCC |
| 3C_C1_B2 | GTGTAATCCCGTGAAGGACTGT |
| 3C_C1_B3 | GGAGCTCTCCAGGTAGGTGA |
| 3C_C1_B4 | TGCTGCTTCTTGCTGGTCAT |
| 3C_C1_B5 | CTTGCCACAAAGCTCACACC |
| 3C_C1_B6 | AACTACCTGGCACACCTTCG |
| 3C_C1_B7 | GGCATTTCGACATTCCTTTCACT |
| C1_B1F | GAGTGCTGGGATTAAAGGCG |
| C1_B1R | TACAAGAACCCACCACCCTC |
| C1_B2F | TTGAGCCAGTACCCATGCAT |
| C1_B2R | GGGATGCCTGTGTCTACTCT |
| C1_B3F | CTTGGCTCCATCCTGTCAGA |
| C1_B3R | TGCCCAGGACACCTTAAACT |
| C1_B4F | ACAGACAGGAAAACATGGTGC |
| C1_B4R | GGAAAGGCTGGGTCACTACT |
| C1_B5F | GCTAGAATCGTGGATGGAAGC |
| C1_B5R | AGACACGGCTGAAGTTCTGA |
| C1_B6F | GAGTTTGCGCTGCTGTGATA |
| C1_B6R | AACGCAAAGCCAAAGGAGTT |
| C1_B7F | AAACACCACGAGAAGCAACC |
| C1_B7R | TGGAGCTCACAACTGTCTGT |
| 3C_C5_B1 | CTCCTTAAGTTGCCCAGGCT |
| 3C_C5_B2 | ATTTCTTGTGTGCATGTCTTCAGT |
| 3C_C5_B3 | CTGGGCATTCTGTCCCACAT |
| 3C_C5_B4 | TGTGTGTACCCTATAGCTGTCC |
| 3C_C5_B5 | AGAACTCTGGCTGCTCTTCC |
| 3C_C5_B6 | TGCTGTCACCTCTCTCCTCA |
| 3C_C5_B7 | TTCTAGTGTGTGTGAGGCCC |
| C5_B1F | TCCTGTGTTTGCCTCCATCT |
| C5_B1R | CATCAGCTCTTTCGAAGCCC |
| C5_B2F | TGATGTCCTCTTCTGGGGTG |
| C5_B2R | GCTCCATGTGCATTTCCCTT |
| C5_B3F | TGCTTTCTGTGGCTCTCTGA |
| C5_B3R | AGGCTAAACAGGTCTGCAGT |
| C5_B4F | GCAGTGCCATTGTCCTTTCA |

| | |
|---|---|
| C5_B4R | AGGAGTCCACAGTTGCAAGA |
| C5_B5F | ATTTGGATGAGGGCGGAAGA |
| C5_B5R | GTTAAGCAGGTGGTGTGGTG |
| C5_B6F | AGAACTAGCTCTGCAGACCA |
| C5_B6R | CCCCGTTTAAAGATGAGCCG |
| C5_B7F | GCATAGCATCACAGAACTTTCCT |
| C5_B7R | ATCTTGCCAAATGAGGTGCA |
| 3C_C10_B1 | ATTACATGCACAAGGCCCTACAT |
| 3C_C10_B2 | TGTGTTTCTGTTCGCCTCCA |
| 3C_C10_B3 | CAGGTCCCTAGCACAACAGA |
| 3C_C10_B4 | CGTAGCGACGTCAGATGAGC |
| 3C_C10_B5 | TGTGTGTAGTTGTTTTGACTGC |
| 3C_C10_B6 | GGTTGCCAGCACTCAAATCA |
| 3C_C10_B7 | TCCTAGCACCTACATGGAAGC |
| C10_B1F | GCACACACAAAGGTCTGGAG |
| C10_B1R | CCTTGCAGTGGGAAATGGAA |
| C10_B2F | ACCCCTAAGCTCTCTCTCCA |
| C10_B2R | TCCTCAGCCCTTACACACAG |
| C10_B3F | ACCTTTGTGTCTGCTGAACA |
| C10_B3R | GGGTCACTTTGATGTTGCGT |
| C10_B4F | AGGGTGACGTGTGGATTAGA |
| C10_B4R | GCAATCTCTCCAGCACCCTA |
| C10_B5F | GCACCTGTGTTTTCCTTGCT |
| C10_B5R | ACCCACATGTCAGCTCACAA |
| C10_B6F | TTCTGACAAGGCCTGGAGAG |
| C10_B6R | CACAGAGGACAGAGAGGGC |
| C10_B7F | TCTGAGGGCTGAAGAGATGTC |
| C10_B7R | GCAAGGCTATTTCATACATGCT |
| ECC3_3C B1 | CTTAGCTGGGACAGTCGTGG |
| ECC3_3C B2 | CTCTTACAGCACTGGGGTCG |
| ECC3_B1F | GGAGACTCGTGGGATGCTTA |
| ECC3_B1R | TCACACACACTGATGCTTAACA |
| ECC3_B2F | CCAAGAAAGGTAAGGCTGCC |
| ECC3_B2R | GTGTAGAGGCCAGGAGTCTC |
| GAPDH_conF | CATCAACGGGAAGCCCATCA |
| GAPDH_conR | CTTTCCGGCCACTTACCC |
